# Inertial interface cavitation creates complex, flow-like structures within a soft solid

**DOI:** 10.1101/2025.02.16.638549

**Authors:** Jin Yang, Alexander McGhee, Zixiang Tong, Griffin Radtke, Mauro Rodriguez, Christian Franck

**Author notes:** J.Y. and A.M. contributed equally to this work.

## Abstract

**Background:** Inertial cavitation near soft material interfaces generates highly asymmetric bubble dynamics, intense stress localization, and complex fluid–structure interactions. However, the subsurface deformation fields within soft solids remain poorly resolved due to limitations in ultrafast, full-field measurement techniques.

**Objective:** This study aims to quantify the spatiotemporal deformation of soft hydrogels during laser-induced inertial cavitation near a gel–water interface.

**Methods:** We integrate single-pulse laser-induced inertial cavitation, an embedded internal Digital Image Correlation (DIC) speckle patterning method, and DIC to resolve *in situ*, full-field subsurface kinematics at 1-2 million frames per second.

**Results:** Among all the tested different non-dimensional stand-off distances, four distinct cavitation–interface interaction regimes are identified, spanning symmetric bulk-like oscillations to strongly asymmetric collapses accompanied by interface indentation, jet reversal, and bubble penetration. Full-field measurements reveal stagnation points, vortex-like deformation patterns, and large localized strains that depend on the non-dimensional bubble stand-off distance.

**Conclusions:** This work establishes an experimental framework for quantifying inertial cavitation dynamics near a compliant gel–water interface. Using ultrafast imaging and DIC, we captured full-field deformation, strain localization, and jet formation across a wide range of stand-off distances near gel-water interfaces.

## 1 Introduction

Inertial cavitation, the rapid expansion and violent collapse of a vapor or gas bubble, plays a central role in a wide range of physical, biological, and engineering processes, from the prey-stunning capabilities of the snapping shrimp [1, 2] to erosion on the propeller blades of naval vessels [3]. In the fluid mechanics community, the dynamics of bubble collapse have been extensively characterized and are known to generate high pressures, shock waves, and microjets capable of producing erosion or facilitating targeted material removal [4, 5]. While cavitation has a long history of scientific study in water and viscous liquids, cavitation in soft matter with complex behavior, i.e., non-Newtonian liquids and viscoelastic soft solids, has recently begun to blossom as a new scientific area of research due to recent significant innovation in experimental, theoretical, and numerical capabilities [6–8].

Classic treatments of bubble dynamics in water, including the Rayleigh-Plesset framework [9, 10] and its extensions [11–21], have provided valuable insight into single-bubble behavior under spherical or weakly asymmetric conditions. However, cavitation in soft viscoelastic solids or along interfaces where fluid and compliant materials interact provides a treasure trove of complex physics [22–27]. That is, a dynamically oscillating bubble in a soft solid is in constant competition with non-equilibrium thermodynamics, large deformation viscoelastic dynamics, and various types of fracture or failure mechanisms [28–32] depending on the magnitude or intensity of oscillation or collapse. This has spawned an exciting new field of soft matter cavitation, which at present has primarily focused on understanding, resolving, and predicting inertial cavitation in bulk soft materials far away from surfaces or impedance-changing interfaces.

However, in many biologically and clinically relevant applications, cavitation is often encountered or purposefully produced along material interfaces, particularly along soft matter-liquid interfaces. Some example applications include cancer cell removal [33, 34], targeted drug delivery [35, 36], cataract removal [37], artificial heart valve maintenance [38], histotripsy [39], noninvasive ocular surgeries [40], and potentially even blast-related traumatic brain injuries [41–43]. Whether cavitation is purposefully generated or an unwanted collateral side effect, one generally seeks to understand the extent of the impact on the surrounding tissue. Thus, being able to quantitatively describe, correctly model (by identifying all pertinent physics) and predict the intricate dynamics of cavitation along these interfaces is paramount to the successful outcome in many of the aforementioned applications.

A central challenge in studying cavitation near soft interfaces is the lack of experimental techniques capable of resolving the subsurface deformation fields with sufficient spatial and temporal resolution. Conventional surface-based methods cannot capture internal strain fields, while traditional imaging approaches lack the frame rates needed to observe transient cavitation dynamics. As a result, many aspects of the fluid–structure interaction remain poorly understood, including how interfacial asymmetry influences bubble morphology, stress localization, jet development, and permanent material deformation.

To address these gaps, this work combines three experimental approaches, including laser-induced inertial cavitation [20, 44–46], Digital Image Correlation (DIC) [47, 48], and the embedded DIC speckle patterning technique [49–51]. This integrated approach enables, for the first time, quantitative measurement of full-field subsurface kinematics in soft solids during interfacial cavitation. By systematically varying the non-dimensional standoff distance between the bubble nucleation site and the interface, we identify distinct regimes of bubble–interface interaction and characterize the associated deformation mechanisms.

The experimental observations presented in this study provide new insight into the mechanics of cavitation near soft interfaces, revealing previously unresolved deformation structures and identifying conditions under which bubble behavior transitions between symmetric, asymmetric, and interface-penetrating collapse modes. These findings improve our understanding of cavitation-driven loading in soft matter and establish a foundation for predictive modeling in applications.

## 2 Materials and Methods

### 2.1 Preparation of gelatin hydrogels with embedded speckle planes

Gelatin hydrogels with an average diameter of 14 mm and an average height of 4 mm were prepared with an internal speckle plane following our previously developed experimental protocols [49–51]. Briefly, porcine gelatin (gel strength 300, Type A; Sigma-Aldrich) was dissolved at three mass concentrations (6 wt%, 10 wt%, and 14 wt%) in deionized (DI) water. The solution was heated and mixed until fully homogeneous, then cast into a mold or glass-bottom dish to form a uniform layer. The gel was first formed with a flat surface after gelation, which was exposed and lightly dried to the extent that the surface could support toner particles without significant dehydration. A thin layer of toner was then deposited onto the surface using a high-pressure air stream to achieve a high-contrast, randomly distributed speckle pattern suitable for DIC analysis.

To embed the speckle layer, a second aliquot of molten gelatin, heated to slightly above the melting temperature of the gel (i.e., about 35^◦^C for 6-14% gelatin gels [52]), was poured over the speckled surface to encapsulate the particles within the bulk. The sample was rapidly cooled in an ice bath to arrest further melting and minimize speckle motion. Finally, the assembled hydrogel was stored at room temperature for about one hour to fuse the layers and eliminate weak interfaces.

Following annealing, all hydrogels were equilibrated in DI water at 4^◦^C for at least 24 h before testing to ensure swelling equilibrium, thereby eliminating time-dependent swelling effects during cavitation experiments. The gel-water interface was subsequently created after equilibration (as schematically illustrated ins Fig. 1(a)), such that no additional swelling-induced evolution of the interface occurred during measurements.

**Fig. 1:**
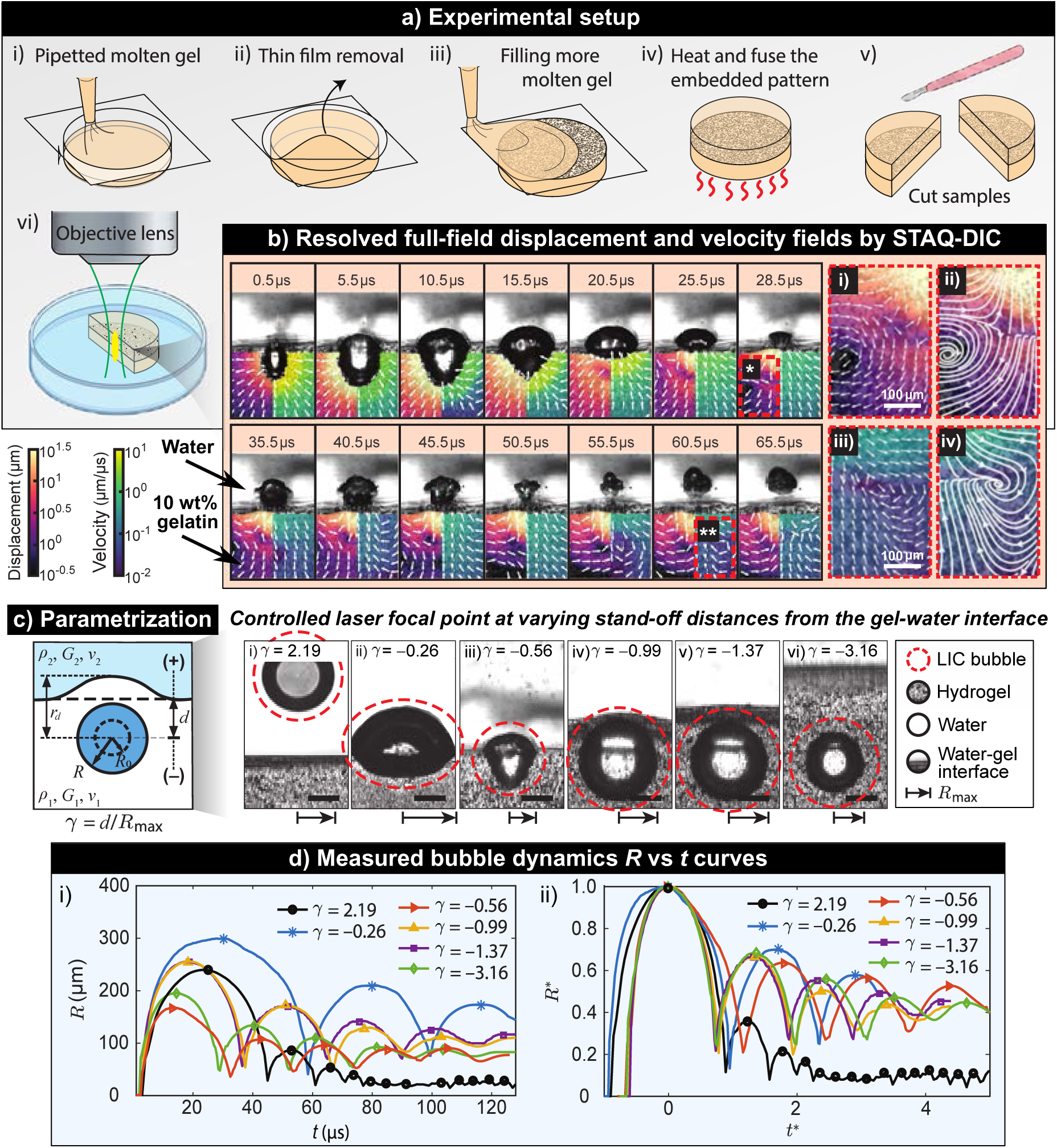
Summary of the experimental setup for the laser-induced inertial cavitation (LIC) experiments at the gel-water interface. (a) Experimental procedures for preparing the embedded speckle pattern in a hydrogel sample. (b) Full-field displacement and velocity fields were resolved using the STAQ-DIC method for a gelatin-water interface under LIC. Selected frames show the evolution of deformation and velocity streamlines in areas of interest (“★” and “★★” regions correspond to (i,ii) and (iii,iv), respectively.) (c) Schematic parametrization of bubble dynamics at varying standoff distances (*γ*) from the gel-water interface. Controlled laser focus positions demonstrate bubble deformation for different *γ* values. Scale bar is 250 µm. (d) Measured bubble dynamics showing bubble radius (*R*) versus time (*t*) in dimensional (i) and normalized (ii) forms, revealing the effects of standoff distance (*γ*) on bubble dynamics.

**Fig. 2:**
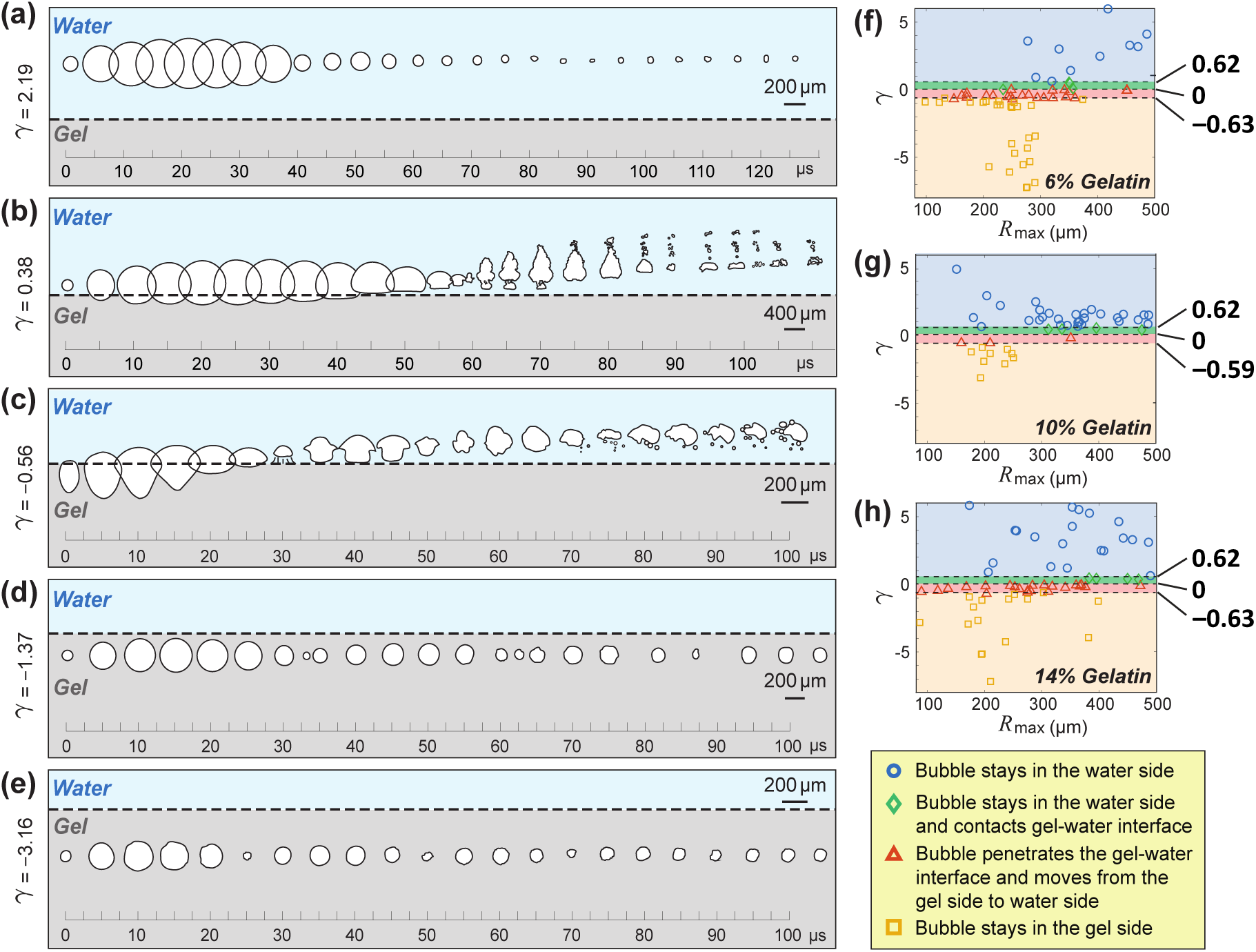
Ensemble-averaged observations of bubble dynamics near a gelatin-water interface, categorized by the non-dimensional stand-off distance *γ* = *d/R*_max_. (a-e) Ensemble-averaged, representative time-lapse morphologies of bubble dynamics during laser-induced cavitation experiments at various *γ* values. (f-h) A summary of the experimentally observed bubble behaviors as a function of their respective maximum radii *R*_max_.

The embedded speckle pattern, formed by micronscale toner particles, is mechanically trapped within the polymer network during gelation and annealing. As a result, once equilibrium swelling is reached, the spatial distribution and stability of the speckle pattern remain unchanged. This is further supported by the high-quality DIC correlation and the agreement between measured displacement fields and the theoretical radial kinematics, indicating negligible relative motion between the particles and the surrounding matrix during experiments.

### 2.2 Quasi-static and high-rate shear modulus measurements

The quasi-static, ground-state shear modulus of each gel was determined using an ARES-G2 rheometer with a 25 mm stainless steel plate (TA Instruments, DE, USA). A strain sweep (0.01%–1% strain at 2 rad/s) was first performed to identify the linear viscoelastic regime for each gel. A strain amplitude of 0.5%, well within the linear regime, was then used for subsequent frequency sweeps.

Frequency sweeps were conducted at 0.5% strain over an angular frequency range of 0.01–10 Hz. Multiple sweeps were acquired to ensure repeatability and minimize Mullins-type effects. The ground-state shear modulus *G*_∞_ was obtained by extrapolating the storage modulus to the low-frequency limit, *f* → 0. The resulting values were *G*_∞_ = 0.74 ± 0.02 kPa, 3.08 ± 0.01 kPa, and 6.71 ± 0.03 kPa for 6 wt%, 10 wt%, and 14 wt% hydrogels, respectively.

To characterize the viscoelastic response at high strain rates relevant to inertial cavitation, Inertial Microcavitation Rheometry (IMR) was employed as described in our earlier work [20, 44–46, 51]. In brief, single cavitation events were generated in each hydrogel and the resulting bubble radius–time histories were fit using a neo-Hookean Kelvin–Voigt visco-hyperelastic model. The dynamic shear modulus *G* and viscosity *µ* extracted from these fits are summarized in Table 1. For the three gelatin concentrations (6%, 10%, 14%), the measured high-rate shear moduli were *G* = 12.78 ± 3.69 kPa, 29.85 ± 6.20 kPa, and 101.50 ± 4.01 kPa, respectively, with corresponding viscosities as 0.027 ± 0.022 Pa·s, 0.055 ± 0.061 Pa·s, and 0.166 ± 0.208 Pa·s, respectively.

**Table 1:** Summary of material properties of used gelatin hydrogels: quasi-static shear modulus G_∞_, high-rate shear modulus *G*, and viscosity *µ* for the gelatin hydrogels used in this study.

| Concentration [%] | $G_\infty$ [kPa] | $G$ [kPa] | $\mu$ [Pa·s] |
| --- | --- | --- | --- |
| 6 (w/w) | $0.74 \pm 0.02$ | $12.78 \pm 3.69$ | $0.027 \pm 0.022$ |
| 10 (w/w) | $3.08 \pm 0.01$ | $29.85 \pm 6.20$ | $0.055 \pm 0.061$ |
| 14 (w/w) | $6.71 \pm 0.03$ | $101.50 \pm 4.01$ | $0.166 \pm 0.208$ |

### 2.3 Laser-induced inertial cavitation and ultra-high-speed imaging

Here, we utilized the same laser-induced inertial cavitation (LIC) setup as previously described in [20, 44, 45, 51]. Briefly, a single Q-switched Nd:YAG laser pulse (pulse duration ∼3–5 ns, wavelength 532 nm) was directed through the back port of an inverted microscope and focused into the hydrogel by a microscope objective (typically 10×/0.3 NA or 20×/0.5 NA). The objective determined the axial location of the nucleation site, while the lateral position of the focal spot relative to the interface was controlled via a motorized microscope stage (see Fig. 1(a:vi)).

In our experiments, various combinations of maximum bubble radius and distance from the interface were tested. As shown in Fig. 1(c), for strongly non-spherical bubbles, an effective radius is commonly defined based on the projected bubble geometry, providing a practical length scale for normalization while not fully capturing shape asymmetry [23, 53, 54]. Here, the effective radius is approximated as half of the maximum projected bubble width.

To normalize these parameters, we utilize the same dimensionless variable (*γ* = *d/R*_max_) as first introduced by Brujan et al. [23, 24] (see Fig. 1(c)), where the bubble standoff distance from the interface is denoted by *d*, and the maximum bubble radius by *R*_max_. We use positive and negative values of *γ* that correspond to initial cavitation nucleation sites in either the water (positive *γ* value) or gel (negative *γ* value), respectively.

The laser-induced cavitation dynamics and subsurface deformation fields were recorded using a Shimadzu HPV-X2 ultra-high-speed camera (Shimadzu Corporation, Kyoto, Japan) operating at 1–2 million frames per second. Each frame was illuminated by a synchronized pulsed laser light source (SILUX640, Shimadzu Corporation) with a pulse duration of approximately 20 ns. The illumination was coupled into the microscope’s epi-illumination path to provide uniform, high-intensity lighting over the field of view. Each recording consisted of 256 frames, including several pre-trigger frames prior to bubble nucleation and the subsequent expansion–collapse cycles. A representative sequence is shown in Fig. 1(b). The bubble radius *R*(*t*) at each time point was extracted from the images using Taubin’s circle fitting algorithm [55]. These data were used both to construct radius–time curves (e.g., Fig. 1(d:i)) and to determine the maximum bubble radius *R*_max_ for each LIC event. Figure 1(d:ii) presents the normalized bubble radius *R*^∗^ versus normalized time *t*^∗^ for various nondimensional stand-off distances *γ*. Non-dimensional quantities are constructed as *R*^∗^ = *R/R*_max_ and *t*^∗^ = *tU_c_/R*_max_, where *U_c_* is the characteristic acoustic velocity which is further defined as 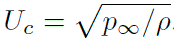.

### 2.4 SpatioTemporally Adaptive Quadtree Mesh Digital Image Correlation

Full-field kinematic deformation measurements in the gelatin were obtained using the SpatioTemporally Adaptive Quadtree mesh Digital Image Correlation (STAQ-DIC) method [48]. Each high-speed frame acquired during an LIC event was compared to the undeformed reference image taken prior to cavitation. The internal speckle pattern provided by the embedded toner particles enabled robust correlation and high spatial resolution within the speckled plane.

STAQ-DIC employs an adaptive quadtree mesh constructed from a binary mask that distinguishes the bubble region from the surrounding material [48, 51]. The bubble silhouette in each frame was segmented to generate a binary mask, and the quadtree mesh was refined near the bubble wall to better resolve large displacement gradients and evolving interface geometry. The coarsest DIC element size was on the order of 8 pixels ×8 pixels, and the finest elements were 2 pixels ×2 pixels.

To improve tracking robustness throughout the cavitation cycle, different DIC tracking strategies were adopted during post-processing. During the early stages of bubble evolution (from t = 0 to the vertical dashed lines in the kymographs shown in Fig. 7(b-d)), incremental frame-to-frame tracking was employed to accurately capture the rapidly evolving deformation field. Incremental displacement fields were first obtained between successive frames in Cartesian coordinates using square subsets centered on each mesh node (DIC subset size is 40 pixels ×40 pixels). Subsets intersecting the bubble region were automatically segmented so that only the material portion (outside the bubble) contributed to the correlation. Incremental displacements were then integrated in time using an accumulation scheme to construct cumulative displacements, following the strategy previously validated for LIC problems [51]. After the time points indicated by the vertical dashed lines in Fig. 7(b-d) and for the entire image sequence of Fig. 7(a), cumulative tracking relative to the undeformed reference configuration was employed to quantify the long-term residual deformation and stress evolution within the hydrogel.

The resulting cumulative displacement fields were then transformed into a polar coordinate system centered at the bubble location. Spatial gradients were evaluated using a local plane-fitting approach with a neighborhood size of 24 pixels ×24 pixels, enabling the computation of full-field finite strain measures, including radial and z-direction strains., as well as derived quantities such as velocity and acceleration fields from temporal finite differences. Representative displacement and velocity streamline fields are shown in Fig. 1(b), and additional examples for different *γ* regimes are provided in Figs. 3–6.

**Fig. 3:**
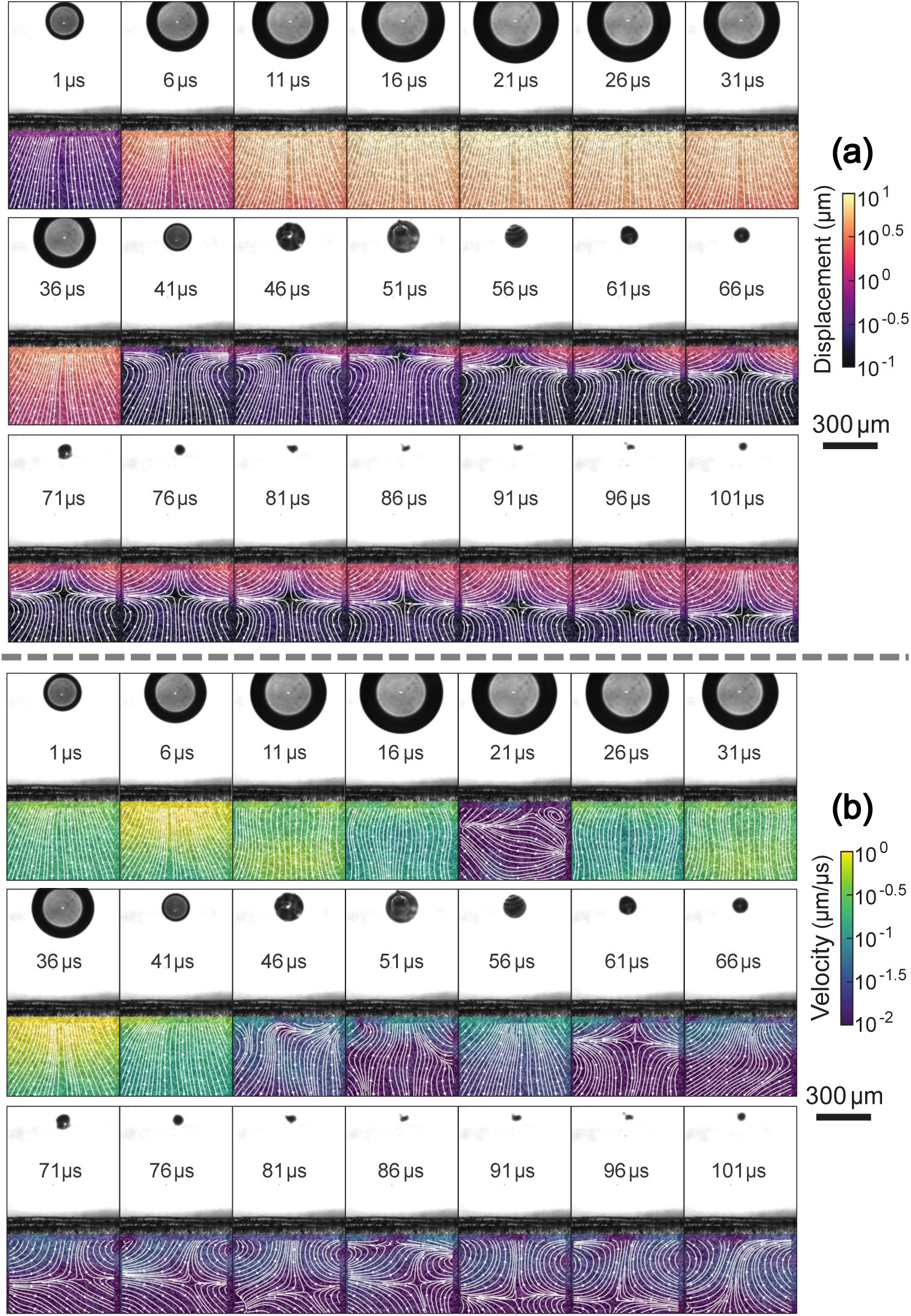
Experimentally-measured, time-resolved analysis of the displacement (a) and velocity (b) field streamlines during laser-induced cavitation near a 10% gelatin gel-water interface with a non-dimensional stand-off distance of *γ* = 2.19.

**Fig. 4:**
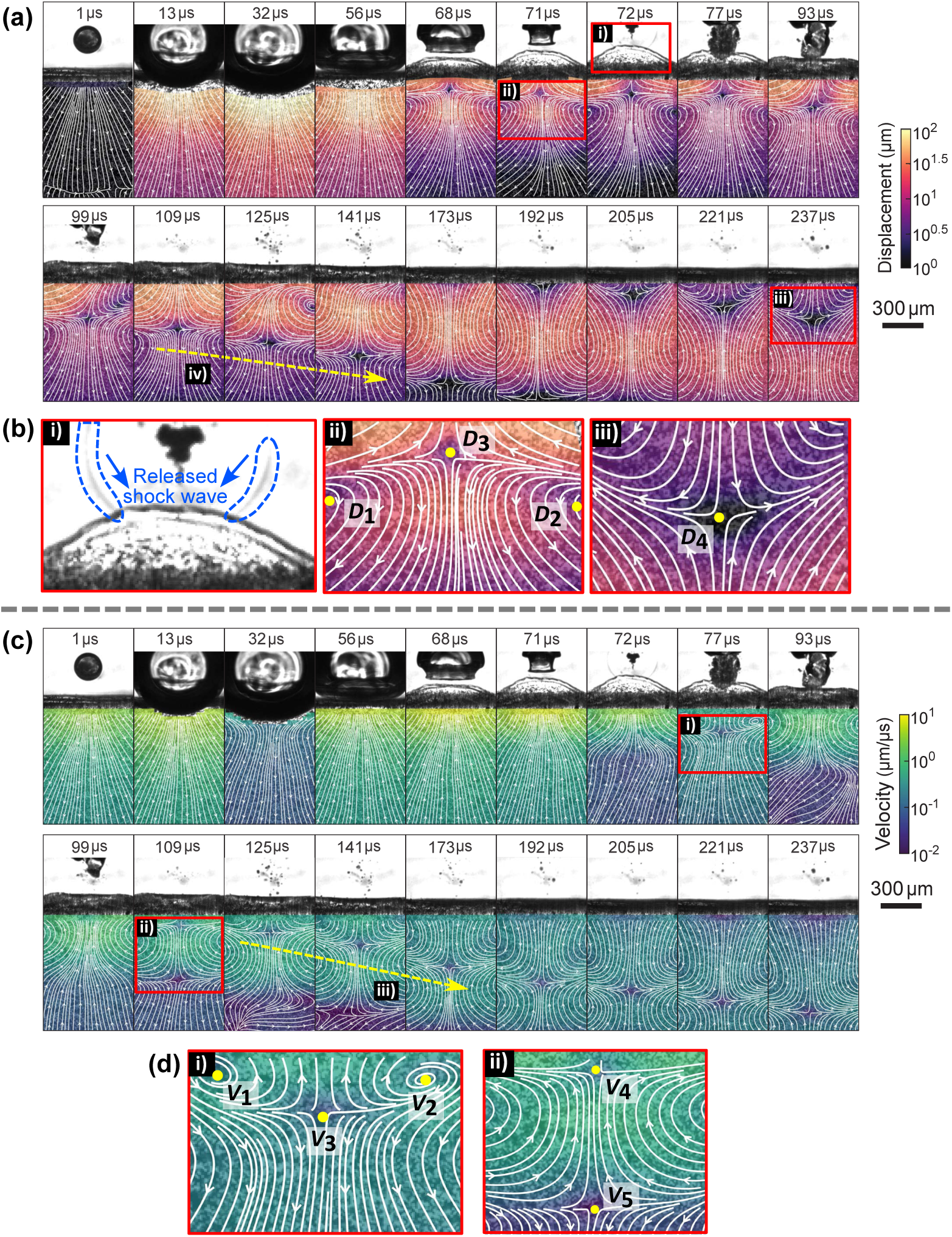
Experimentally-measured, time-resolved analysis of the displacement (a) and velocity (c) field streamlines during laser-induced cavitation near a 10 wt% gelatin gel-water interface with a non-dimensional stand-off distance of *γ* = 0.38. (a:i-iii) are enlarged and shown in (b). Two dashed yellow lines marked in (a:iv) and (c:iii) illustrate the shear stress wave propagation beneath the gel-water inter-face. (b:i) A shock wave is released upon bubble collapse. (b:ii,iii) show detailed views of the displacement fields at 71 µs and 237 µs. Identified stagnation points (*D*_3_, *D*_4_) and vortex pairs (*D*_1_, *D*_2_) are indicative of complex displacement fields resulting from the bubble’s collapse. (c:i,ii) are enlarged and shown in (d). (d) shows detailed views of the velocity fields at 77 µs and 109 µs. Identified stagnation points (*V*_3_, *V*_4_, *V*_5_) and vortex pairs (*V*_1_, *V*_2_) illustrate the complex velocity field evolution induced by the collapsing bubble.

**Fig. 5:**
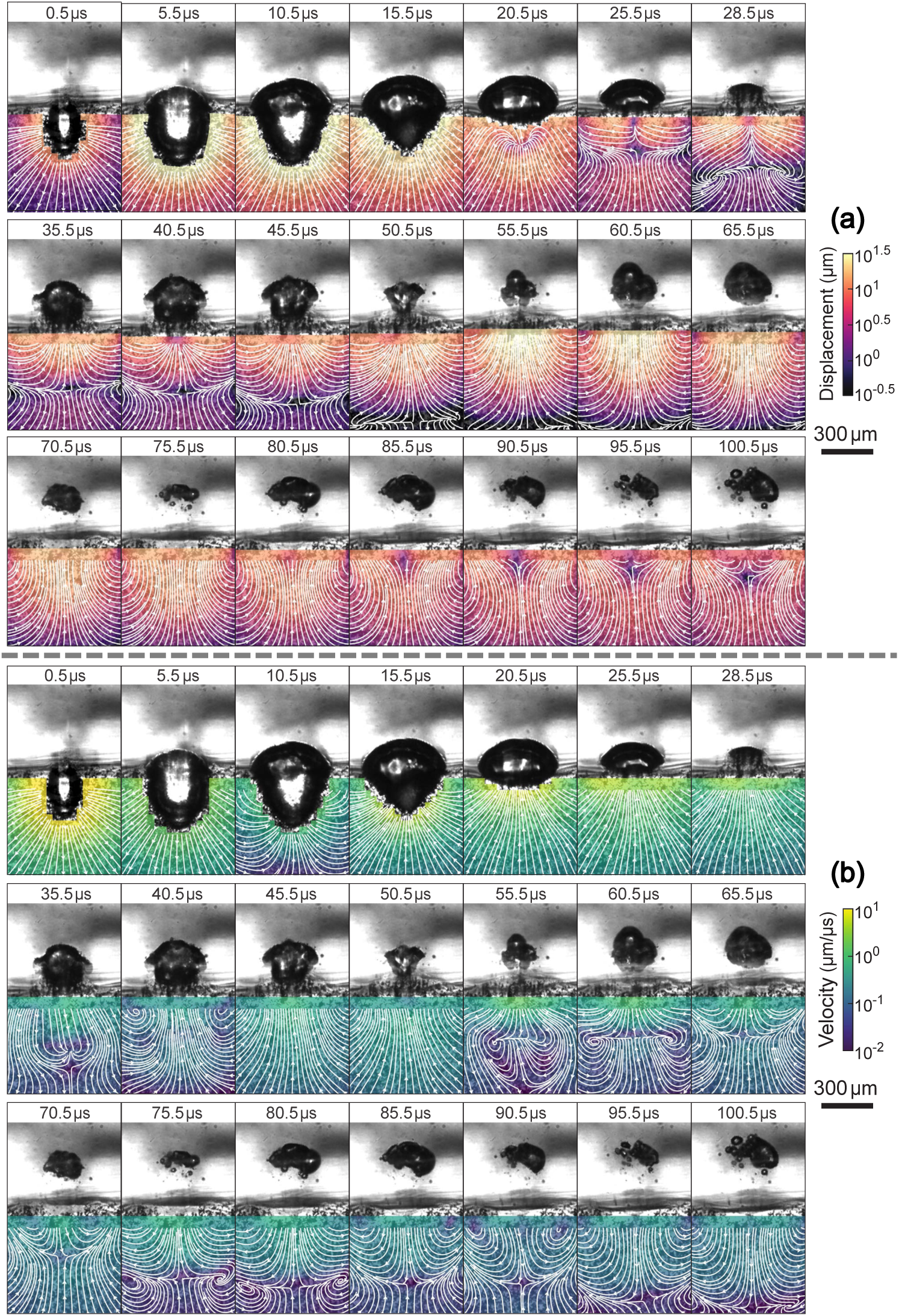
Experimentally-measured, time-resolved analysis of the displacement (a) and velocity (b) field streamlines during laser-induced cavitation near a 10% gelatin gel-water interface with a non-dimensional stand-off distance of *γ* = −0.56.

**Fig. 6:**
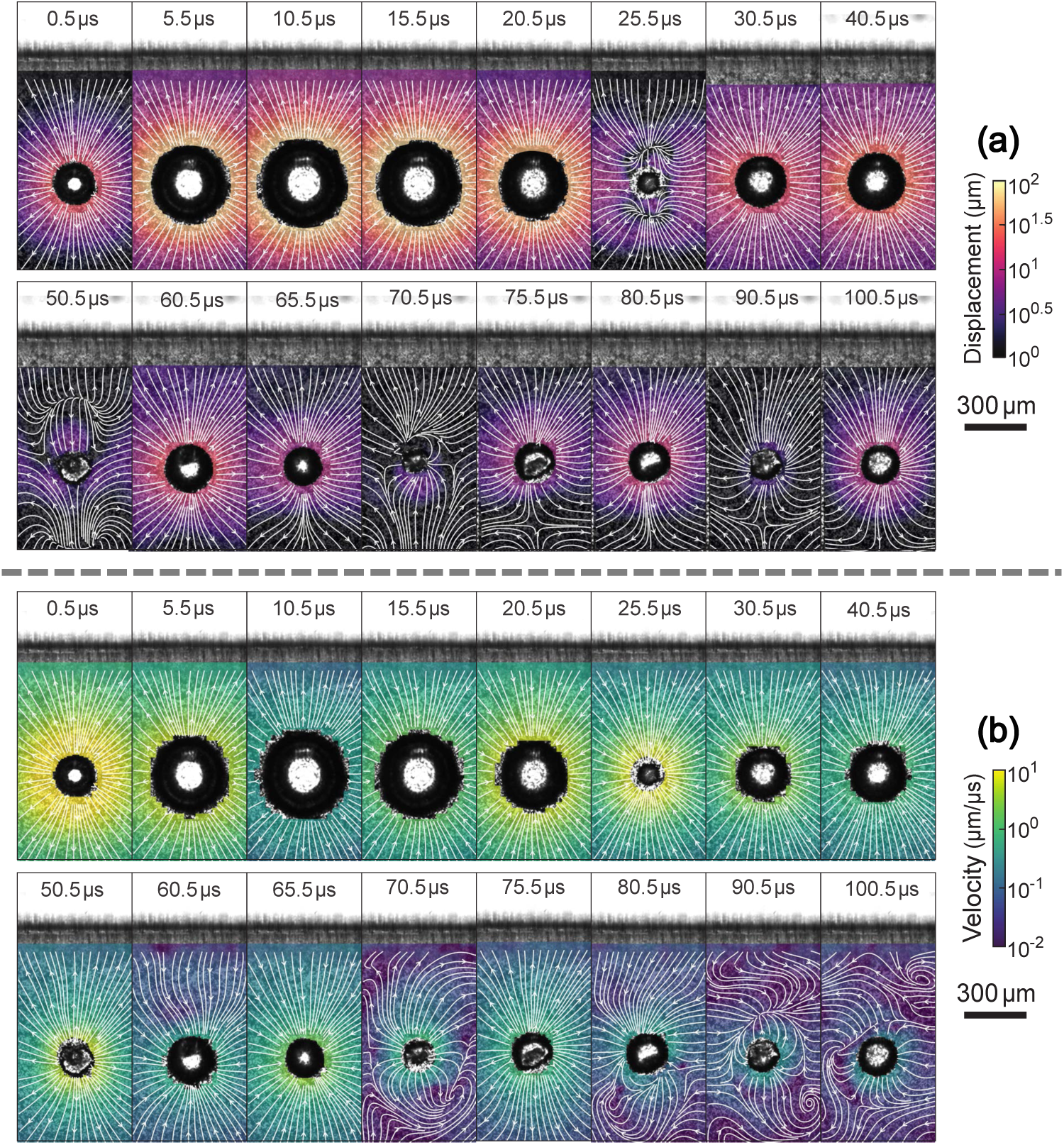
Experimentally-measured, time-resolved analysis of the displacement (a) and velocity (b) field streamlines during laser-induced cavitation near a 10% gelatin gel-water interface with a non-dimensional stand-off distance of γ = −3.16.

**Fig. 7:**
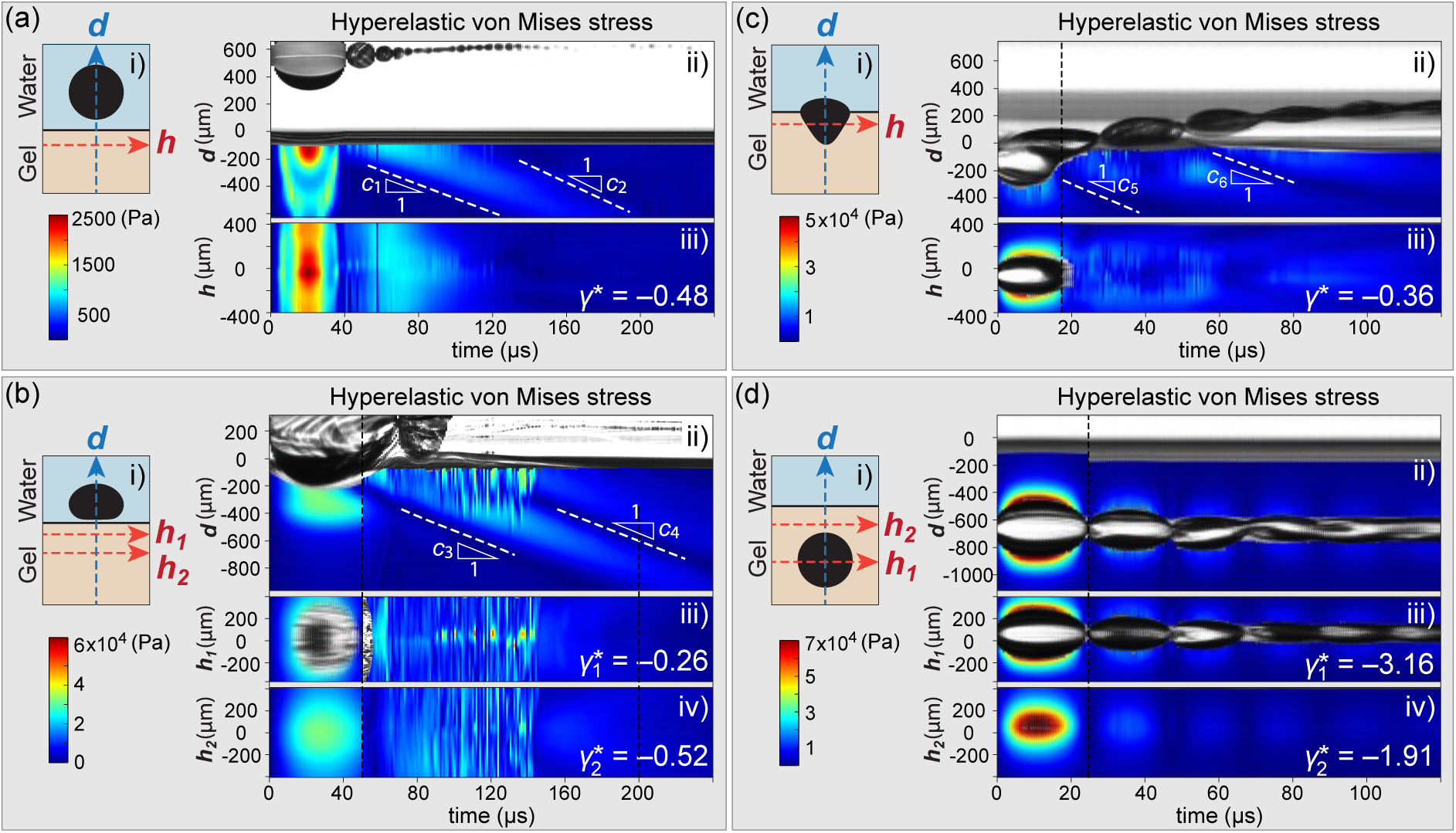
Experimentally measured kymographs of hyperelastic von Mises stress generated by laser-induced inertial cavitation dynamics at a 10% gelatin gel-water interface as a function of non-dimensional stand-off distance *γ*: a) *γ* = 2.19, b) *γ* = 0.38, c) *γ* = −0.56, and d) *γ* = −3.16. In each subfigure, i) shows the non-dimensional stand-off distance; ii-iii) or (ii-iv) present time-resolved stress distribution along *d* and *h* axes. Measured wave propagation speeds: *c*_1_ = 5.0 m/s, *c*_2_ = 6.1 m/s, *c*_3_ = 9.2 m/s, *c*_4_ = 5.5 m/s, *c*_5_ = 7.5 m/s, *c*_6_ = 9.1 m/s. The vertical dashed lines in (b-d) indicate the transition between two DIC tracking strategies used during post-processing. Prior to the dashed line, incremental frame-to-frame tracking was employed to improve robustness during rapid bubble collapse and rebound. After the dashed line, cumulative tracking relative to the undeformed reference configuration was used to quantify long-term residual deformation and stress evolution. (a) was tracked using cumulative tracking mode only.

### 2.5 Stress calculation from DIC results

To estimate the local stress fields associated with the experimentally measured deformation patterns, we post-process the full-field DIC displacement measurements using a finite-deformation viscohyperelastic constitutive framework. In our experiments, we assume the displacements are axisymmetric, that *θ*-direction displacement component *u_θ_* = 0. Based on our experimental observations, this approximation holds for the majority of all measured bubble life cycles. Both *u_r_* and *u_z_* displacement components in the gelatin hydrogels can be directly measured from DIC results. We consider the deformation in cylindrical coordinates from the undeformed reference configuration (*R, Θ, Z*) to the deformed configuration (*r, θ, z*), where the deformation mapping is therefore expressed as

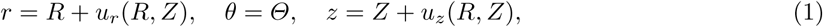

where u*_r_* and u*_z_* are r- and z-direction displacement components, respectively.

Under the axisymmetric assumption, the deformation gradient tensor in cylindrical coordinates is

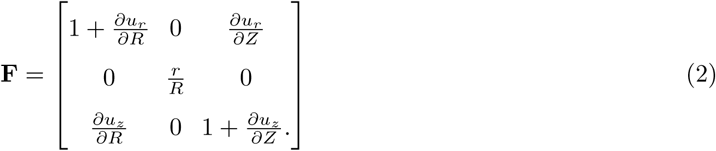

To estimate the stress field, the gelatin hydrogel is modeled as a neo-Hookean Kelvin–Voigt visco-hyperelastic material [20, 44, 56]. The corresponding Cauchy stress tensor, ***σ***, is written as

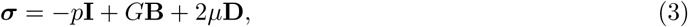

where *p* is the pressure term enforcing incompressibility; **I** is the identity tensor; *G* is the dynamic shear modulus; and *µ* is the effective viscosity. The material parameters *G* and *µ* were previously determined using our IMR framework applied to cavitation in bulk gelatin hydrogels [20, 44–46, 51] (see Table 1). Variable **B** is the left Cauchy–Green deformation tensor that is calculated as

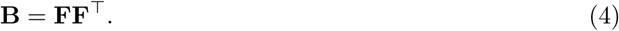

The experimentally measured displacement fields are further differentiated in time to compute the velocity field components,

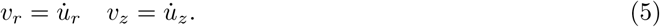

The corresponding spatial velocity gradient can be calculated as

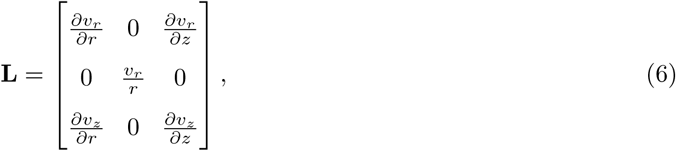

from which the rate-of-deformation tensor is calculated as

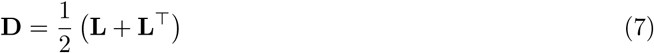

To quantify the intensity of the local stress concentration, we further compute the von Mises equivalent stress from the deviatoric component of the Cauchy stress tensor. The deviatoric stress tensor, **s**, is defined

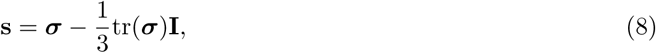

and the corresponding von Mises stress is

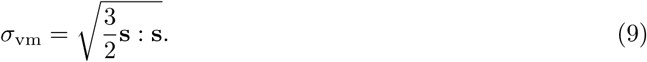

For the present axisymmetric formulation, the von Mises stress can be written explicitly as

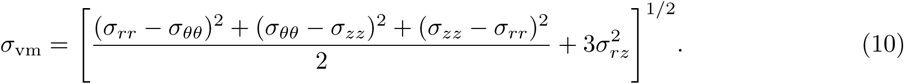

Using the experimentally measured full-field displacement and velocity data together with the constitutive relations above, we estimate the spatiotemporally evolving hyperelastic and viscoelastic stress fields generated during interfacial cavitation for different stand-off distances *γ*. These stress fields provide quantitative insight into the localized stress concentration, wave propagation, and residual deformation patterns induced by strongly nonlinear cavitation dynamics near the gel–water interface.

## 3 Results and Discussion

### 3.1 Classification of Bubble–Interface Interaction Regimes

A systematic series of cavitation experiments was conducted for bubbles nucleated at different stand-off distances from the gel–water interface. The non-dimensional stand-off parameter *γ* = *d/R*_max_ was used to classify the interaction regimes. For large positive stand-off distances (*γ* ≫ 1), the cavitation bubble underwent nearly spherical expansion and collapse, similar to classical dynamics in an unbounded medium [57, 58]. As the bubble was positioned closer to the interface (*γ <* 1), a rebound jet directed toward the interface emerged [23, 24]. When cavitation was nucleated on the gel side, i.e., negative *γ* values, but close to the gel–water interface, the collapse became increasingly asymmetric, with the jet reversing direction and penetrating into the gel. This systematic variation indicates that *γ* is an effective organizing parameter governing the strength and orientation of asymmetry in cavitation near a compliant boundary.

Contrary to previous studies that focused solely on examining the bubble dynamics in water with regard to a gel-water interface (i.e., positive *γ* values) [23, 24], our research explores both positive and negative values of *γ*. Four qualitatively distinct behaviors were observed, ranging from fully bulk-like spherical collapse to strongly asymmetric, interface-dominated jetting. Figure 2 provides a general summary of our interfacial cavitation observations with Fig. 2(a-e) displaying representative, spatiotemporally varying bubble morphologies, and Fig. 2(f-h) summarizing general observations of bubble movement.

The observed bubble dynamics are clearly sensitive to the dimensionless variable *γ*, leading to particularly complex morphologies for |*γ*| *<* 0.62. We observe that for *γ_c_ < γ <* 0, a bubble would be nucleated on the gel side and then migrate across the interface. From our measurements, the critical stand-off distance for penetration is approximately *γ_c_* ≈ −0.62 (*γ_c_* = −0.63, −0.59, and −0.63 for 6 %, 10 %, and 14 % gelatin hydrogels, respectively). Notably, these values are nearly invariant with concentration, indicating that the critical stand-off distance *γ_c_* is relatively insensitive to changes in gelatin content within the range studied.

While the high-rate shear modulus increases approximately monotonically with concentration [46], the weak dependence of *γ_c_* on concentration indicates that stiffness alone does not control the penetration threshold. Instead, when the bubble crosses the gel-water interface, the material undergoes extremely large, highly localized deformations, accompanied by significant dissipation and likely localized fracture damage. Therefore, the relevant parameter governing *γ_c_*is more closely associated with the material’s resistance to effective dynamic fracture toughness rather than its elastic modulus.

For 0 *< γ <* 0.62, the nucleated bubble stays on the water side and indents the gel-water interface. In the present experiments, for all tested gelatin hydrogels (6–14 wt%), we did not observe bubbles penetrating from the water phase into the gel phase for any positive *γ* values. Even for small positive stand-off distances approaching the interface, bubbles nucleated in the water phase remained confined to the water side throughout the cavitation cycle. Interestingly, previous studies have reported qualitatively different behavior in other soft materials, where sufficiently small but still positive *γ* values can lead to bubble splitting and partial penetration into the compliant material side [54]. A more detailed discussion of this regime and associated jetting dynamics can be found in Ref [54].

Finally, when *γ >* 0.62 or *γ < γ_c_* ≈ −0.62, each of the nucleated bubbles remained in its respective initial phase, irrespective of the gel’s stiffness. That is, a bubble nucleated in water or gelatin did not cross the interface. The resulting bubble dynamics are consistent with previous literature observations of cavitation near an elastic interface or within a bulk hydrogel [17, 19, 59]. As the stand-off distance increases relative to the maximum bubble radius, the asymmetry in the pressure field induced by the interface decays rapidly, and the bubble dynamics approach those of an infinite medium. Consequently, jet formation and interface deformation are suppressed, and the bubble exhibits nearly symmetric expansion and collapse. Within the range of material properties investigated, variations in gel stiffness are small compared to the characteristic pressures generated during cavitation, and therefore do not significantly affect this transition, which is primarily governed by geometric effects.

### 3.2 Measured spatiotemporally evolving full-field deformation fields

To understand the potentially complex soft matter deformation behavior during bubble growth and collapse, we experimentally measured the deformation characteristics within each of the four *γ* regimes. For brevity’s sake, Figs. 3-6 present kinematic time-lapse data of our 10% gelatin gels. While the magnitudes show differences depending on the stiffness of the gel, the general kinematics are similar among all gels, and well represented in Figs. 3-6.

#### 3.2.1 (a) γ > 0.62: Cavitation in the water phase without interface contact

For *γ >* 0.62, cavitation bubbles nucleated and remained on the water side, which was consistently observed across all three different gel stiffness (Fig. 2(a, f-h)). By plotting the displacement and instantaneous velocity field streamlines (see Fig. 3), we observed relatively small deformations with maximum principal strains less than 1.5% across the full cavitation cycle (see Supplementary Materials Fig. S1).

The displacement streamlines reveal a radially symmetric deformation pattern during the bubble’s initial expansion and collapse phase reaching R_max_ at approximately 21 µs (see also Fig. 1(d)) with an associated maximum material displacement close to 10 µm and an accompanying instantaneous velocity field near zero, which is consistent with prior theoretical treatments [20]. As the bubble enters the initial collapse phase (Fig. 3: 21 µs ≲ *t* ≲ 41 µs) we see that the subsurface deformations remain largely radially symmetric until after the first collapse. Furthermore, one can clearly see a reversal in the instantaneous material velocity fields as soon as the bubble enters the collapse phase (Fig. 3: *t* = 26 µs), with the velocity magnitude increasing rapidly as the bubble approaches its first collapse. While one might expect the largest velocities to occur immediately at the collapse point (Fig. 3, *t* = 41 µs), the measured velocities within the gelatin appear slightly smaller than *t* = 36 µs. We attribute this discrepancy primarily to the rapid rebound of the bubble immediately following collapse, during which the radial material velocity reverses direction from inward to outward over a very short timescale. Due to the finite temporal resolution of the high-speed imaging system and the associated DIC post-processing, these rapid velocity reversals are likely partially temporally averaged, leading to an underestimation of the peak instantaneous material velocity near collapse.

Shortly after the first collapse (i.e., Fig. 3(a) at *t* = 46 µs), a stagnation point emerges within the displacement streamline field. This stagnation point marks the transition between outward residual deformation generated during bubble expansion and the newly induced inward motion associated with bubble collapse. Physically, the rapid deceleration and reversal of the surrounding material motion during collapse generate strong shear deformation near the compliant gel–water interface, leading to localized tangential motion and the nucleation of a propagating shear wave packet. Both displacement and velocity streamline topologies further reveal that the collapse-induced flow field is no longer purely radial, but instead contains substantial rotational and shear-dominated motion caused by the asymmetric bubble collapse near the interface. As shown in Fig. 3(b), at later times (*t >* 46 µs), although the bubble oscillation amplitude has substantially decayed, persistent velocity fluctuations and weak rotational structures remain visible near the interface region, suggesting that a portion of the collapse energy continues to redistribute through long-lived viscoelastic relaxation and propagating shear-wave motion within the hydrogel.

We investigated the shear wave speeds, which can be visualized more clearly by constructing kymographs of the von Mises stress field shown in Fig. 7(a), which we compute from the experimentally measured deformation field and by assuming a hyperviscoelastic Kelvin-Voigt model [20] using the material properties from Table 1. Here, for each frame of the DIC results, a 7-pixel wide slice along the middle vertical dashed line is cropped and then concatenated to produce Fig. 7(a:ii). Similarly, a 7-pixel wide slice along the horizontal dashed line (*γ*^∗^ = −0.48) is concatenated to create Fig. 7(a:iii). The cavitation bubble indents the gel, and when it reaches its maximum radius, it produces a peak von Mises stress of approximately 2,500 Pa (Fig. 7(a)). At this point, all strain components remain below 1.5% (see details in Supplementary Material Fig. S2).

Following the first collapse, the material below the interface generates shear deformations propagating at speeds of *c*_1_ = 5.0 m/s and *c*_2_ = 6.1 m/s, respectively (Fig. 7(a), Supplementary Materials Section S1 and Fig. S1). Both shear wave speed values are consistent with our earlier IMR measurements of the underlying dynamic material shear modulus (Table 1) and density [46, 51] (*c_s_* ∼ *G/ρ*). We hypothesize that the initially slower wave speed is due to the existence of significant compressibility effects near the bubble collapse point, which reduce the effective instantaneous shear-wave propagation speed immediately following collapse [44]. Ultimately, the laser-induced bubble dissolves into the surrounding liquid, and the gel material elastically recovers to its original state.

#### 3.2.2 (b) 0 < γ < 0.62: Cavitation in the water phase with interface contac

For cavitation bubbles nucleated within a stand-off distance 0 *< γ <* 0.62 (Fig. 2(b, f-h)), the bubble significantly indents the interface and induces large deformations in the gel, the induced strain field is more than a magnitude larger than what was observed for *γ >* 0.62 (see Fig. 4).

During the initial expansion phase, both the displacement and velocity fields remain radially symmetric, a behavior that is largely maintained until the bubble enters the collapse phase around *t* = 32 µs (Fig. 4). As a result, both the displacement and velocity fields exhibit nearly radial streamline patterns. However, once the bubble approaches the interface and enters the collapse phase (right after *t* = 56 µs in Fig. 4), the pressure field becomes highly asymmetric due to the large impedance mismatch between the water and gelatin phases. This asymmetry produces nonuniform interface acceleration and directional momentum transfer, which substantially alters the surrounding deformation field.

Specifically, during LIC bubble expansion, material points move predominantly outward, whereas bubble collapse induces strong inward motion toward the shrinking cavity. Near the compliant gel–water interface, the collapsing bubble generates a strong pressure gradient that pulls the hydrogel surface upward toward the bubble. However, due to the viscoelastic nature of the gelatin, the deeper subsurface region still retains residual downward motion associated with the earlier expansion phase. As a result, the near-surface material is pulled upward while the deeper region continues to move downward, producing a transition region where the opposing motions balance each other. This competition gives rise to a stagnation point at *t* = 68 µs (Fig. 4(a)). We observe the emergence of slight perturbations in the otherwise radial velocity and displacement fields, which add to the propagating tension-compression stagnation point at *t* = 71 µs (Fig. 4(a:ii) and (b:ii)), featuring both significant radial stretches (area around point *D*_3_) as well as noticeable shear deformations visualized by the rotational cores at points D_1_ and D_2_.

As the bubble collapses (at *t* = 72 µs) we observe large interface deformations with the release of a shock wave (Fig. 4(b:i)), and the generation of significant shear deformations marked by an emergence of recirculation zones, both in the displacement and velocity fields (Fig. 4(c:i-ii) and (d:i) points *V*_1_ and *V*_2_). These patterns are largely driven by shear wave deformations, causing the material to exhibit rotational motion (recirculation). Examining the velocity field underneath the expanding and collapsing bubble cloud at *t* = 77-109 µs in detail (Fig. 4(d:i-ii)), we find the generation of vortex ring-like structures that become significantly stretched and compressed by the oscillating free surface deformations giving rise to a second stagnation point upon reversal of the sub-surface velocity field at 109 µs (see Fig. 4: point *V*_3_ in (d:1) and point *V*_5_ in (d:ii) are the first stagnation point; (d:ii) point *V*_4_ is the second stagnation point). The propagation of shear waves is evident in both the displacement (Fig. 4(a:iv)) and velocity (Fig. 4(c:iii)) streamlines. As before, we experimentally measure the shear wave speeds and find c_3_ = 9.2 m/s and c_4_ = 5.5 m/s after the first and second collapse (see Fig. 7(b) for more details). The propagation speed of the second shear wave, c_4_, is slower than that of the first shear wave, c_3_. We attribute the significant reduction in the shear wave speed to the decrease in the dynamic shear modulus due to accumulated material damage stemming from excessive deformations.

In stark contrast to the radially dominated deformation field for γ > 0.62, even at late time points (i.e., *t >* 125 µs), we also observe significant residual displacement fields persisting after the primary cavitation event, particularly concentrated near the gel–water interface (see Fig. 4(a), *t* = 192–237 µs and a second stagnation point *D*_4_ in Fig. 4(b:iii)). This behavior suggests that the interface region does not fully recover immediately after bubble collapse, but instead retains long-lived deformation associated with viscoelastic relaxation and irreversible energy dissipation. During collapse, the bubble generates strong localized pressure gradients and highly non-uniform deformation fields near the compliant free surface. Part of the deformation energy is elastically recovered during rebound, while another portion is dissipated through viscous damping and potentially localized microstructural damage within the gelatin network. As a result, the interface exhibits residual upward bulging and persistent displacement gradients even after the main bubble oscillations have decayed.

In addition, the non-spherical collapse and subsequent jetting process introduce asymmetric momentum transfer into the surrounding hydrogel, producing complex recirculation-like displacement patterns and delayed relaxation near the interface. The persistence of these residual displacement fields indicates that the interface response is governed not only by instantaneous inertial loading, but also by the coupled effects of viscoelastic relaxation, free-surface deformation, and localized damage accumulation under high-rate cavitation loading.

It is worth noting that the displacement magnitudes during the expansion phase appear larger than those during collapse. This apparent asymmetry is primarily a consequence of the measurement window and the cumulative definition of displacement. In the present experiments, all speckles are embedded below the gel–water interface, allowing the motion of nearly the entire tracked region to be resolved during expansion as the bubble indents into the gel. During collapse, however, material motion partially reverses the earlier deformation, and the largest displacements occur very close to the free surface. These near-surface deformations are not fully captured because they lie outside the DIC post-processing field of view and because the speckle pattern becomes sparse in this thin region.

As a result, the peak displacements during collapse are underestimated. If the measurement window extended above the interface, even larger surface displacements would likely be observed. Therefore, the apparent asymmetry between expansion and collapse phases should be interpreted as a limitation of the measurement rather than a fundamental physical effect.

#### 3.2.3 (c) −0.62 < γ < 0: Cavitation penetration through the interface

As shown in Fig. 2(c, f-h), cavitation occurs within the gel near the interface when −0.62 *< γ <* 0. In this regime, the bubble generally migrates from the gel half into the lower impedance water half, thus penetrating the gel-water interface [60]. Figure 5 presents the experimentally measured displacement and velocity streamline evolution for a cavitation bubble nucleated inside the gelatin phase at a non-dimensional stand-off distance of *γ* = −0.56. Unlike the cases with positive stand-off distances, where the bubble remains primarily in the water phase and only indents the compliant interface, the present case corresponds to an interface-penetrating cavitation event in which the expanding bubble directly interacts with and subsequently traverses the gel–water interface.

During the early expansion phase (*t* ≲ 5.5 µs in Fig. 5), the surrounding gelatin exhibits a predominantly outward radial displacement field, similar to bulk cavitation behavior. However, because the bubble is nucleated close to the compliant free surface, the deformation field rapidly becomes asymmetric. The interface deforms upward toward the water phase, and the streamline patterns reveal strong vertical stretching near the bubble apex. Compared with the *γ >* 0 cases, the displacement field is substantially more localized near the interface, indicating that the free surface strongly modifies the surrounding stress field and breaks the near-spherical symmetry of the deformation.

As the bubble reaches its maximum size (*t* ≈ 10.5 µs in Fig. 5), due to the significant stiffness mismatch between the gel and the water, the upper portion of the bubble protrudes into the water phase while the lower portion remains confined within the gelatin. The bubble’s initial elliptical shape quickly degenerates into an inverse teardrop shape with a semi-spherical dome expanding into the water half. This partial penetration generates highly non-uniform deformation gradients near the interface. In particular, the streamline topology shows that material points near the surface are redirected upward and laterally along the interface rather than purely radially away from the bubble center.

During collapse (15.5 µs ≲ *t* ≲ 28.5 µs in Fig. 5), the velocity field reverses direction and becomes substantially more complex than during expansion. Strong inward motions toward the collapsing cavity develop beneath the interface, while the near-surface region exhibits pronounced lateral flow-like motions and rotational structures. These recirculation-like streamline patterns are consistent with asymmetric collapse dynamics induced by the nearby compliant interface (see Case (b): 0 *< γ <* 0.62). The collapse no longer behaves as a purely radial inertial contraction; instead, the interface geometry redirects momentum tangentially along the surface, generating localized rotational cores and stagnation regions. We observe the generation of a vortex ring-like structure in the displacement streamlines (Fig. 5(a)) at *t* = 25.5-50.5 µs followed by vertical bubble motion deeper into the water half. During collapse, high deformation regions form near the bubble boundary, with shear wave propagation becoming evident as radially outward-moving deformation patterns in the gel, starting at 20.5 µs, as seen in the displacement streamlines in Fig. 5(a). As before, we experimentally measure the shear wave speeds and find *c*_5_ = 7.5 m/s and *c*_6_ = 9.1 m/s after the first and second collapse (see Fig. 7(c) for more details). The time points *t* = 28.5 µs to *t* = 80.5 µs feature a unique interplay between a stagnation point generated from rapid changes in directional velocity due to fast-moving longitudinal waves, which can be clearly visualized by periodic direction changes of the normal velocity component and downward propagating vortex ring-like structures (see Fig. 5(b)).

These cycles of complex expansion and collapse cause significant damage to the gel material, which can be seen in some of the frames when closely inspecting the dynamically evolving interface contour. At later time points (*t* ≳ 85.5 µs) (see Fig. 5(a)), persistent residual displacement and velocity fields remain concentrated near the interface even after the main bubble oscillation has decayed. In contrast to bulk-like cavitation, where the deformation field largely relaxes after rebound, the interface-penetrating case retains long-lived deformation signatures, even after the bubble dissolves in the water. In fact, this mode of cavitation is the most destructive of the four observed scenarios. It results in permanent residual strains reaching approximately 20% and hyperelastic von Mises stresses on the order of *O*(10) kPa near the water-gel interface, as indicated in Fig. 7(c) and Supplementary Materials Fig. S3. Notably, the critical cutoff at *γ* ≈ −0.62 is insensitive to the gel modulus across all tested gelatin concentrations. This insensitivity may be governed by the large impedance mismatch across the interface, which may dominate any stiffness or microstructurally dependent failure modes [61] driving the bubble across the interface.

#### 3.2.4 (d) γ < −0.62: Cavitation in the gel without interface contact

When *γ <* −0.62 (see Fig. 2(d-e, f-h)), the laser-induced inertial bubble nucleates and remains entirely within the gel half. This behavior closely resembles cavitation within an infinite bulk solid [20, 21, 51, 62]. As shown in Fig. 6, and consistent with our prior experimental and analytical treatments of inertial cavitation within a solid, the displacement and velocity fields remain almost entirely radially symmetric throughout the entire cavitation process. Noticeable deviations occur primarily during the bubble collapse and final end cycle stages where both the displacement and velocity magnitudes are small. As the bubble reaches its maximum radius (*t* = 10.5 µs in Fig. 6), the velocity field diminishes significantly, becoming almost negligible. However, during collapse, the velocity field reverses direction, with inward motion dominating as the bubble contracts. This is in contrast to the displacement field, which continues to display outward deformation due to its cumulative nature. Near the first collapse (*t* = 25.5 µs in Fig. 6), the velocity field attains its maximum, consistent with theoretical prediction of an accelerating radial contraction of the bubble, while the displacement field shows two stagnation points in the displacement streamline plot. After the second collapse at 70.5 µs (Fig. 6), rotational motion via the emission of shear wave propagation is noticeable, causing slight deviation in the otherwise radially-symmetric field.

This conclusion is further supported by the experimentally measured kymographs in Fig. 7(d). The generated hyperelastic von Mises stress fields exhibit near-perfect spherical symmetry, as evidenced by the minimal differences between the 7-pixel-wide vertical slice along line *d* and the horizontal slice along line *h*_1_ (γ^∗^ = −3.16). For comparison, another horizontal slice at line *h*_2_ (*γ*^∗∗^ = −1.91) is also visualized, further confirming the observed symmetry.

### 3.3 Influence of stand-off distance on residual strain distribution in gelatin hydrogels

We further quantify the influence of stand-off distance on residual strain distribution in subsurface soft matter and use gelatin 10 wt% as a material model system. Figure 8 summarizes the residual von Mises strain distribution^1^ in a 10 wt% gelatin hydrogel subjected to laser-induced cavitation at varying stand-off distances from the gel-water interface. The pink dashed lines indicate the gel-water boundary. The non-dimensional stand-off distance *γ* is labeled near each image. These results highlight the influence of boundary proximity on the material damage of gelatin hydrogels to cavitation-induced deformation.

**Fig. 8:**
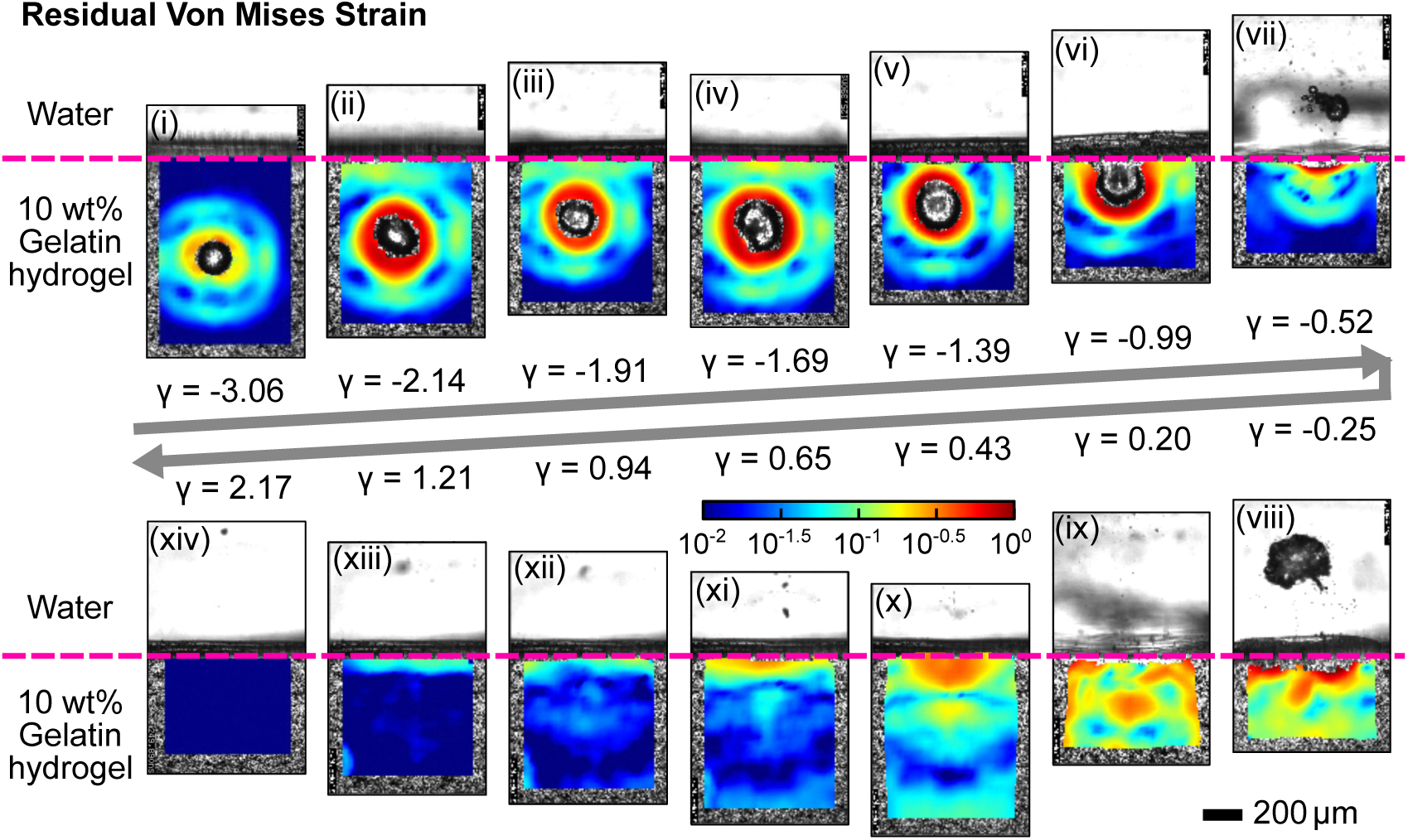
Residual von Mises strain distribution in a 10 wt% gelatin hydrogel subjected to varying stand-off distances to the gel-water interface caused by laser-induced cavitation. Gel-water interface is indicated by pink dashed lines.

The results show that *γ* strongly governs both the magnitude and spatial localization of residual strain. For large stand-off distances (*γ >* 0.62), the residual strain is minimal, consistent with nearly symmetric, bulk-like cavitation dynamics. For *γ <* −0.62, significant residual strains develop near the residual bubble area.

In contrast, for intermediate stand-off distances (|*γ*| < 0.62), significant residual strains develop near the gel-water interface due to strong interface indentation and asymmetric collapse. Particularly in the regime −0.62 *< γ <* 0, where cavitation-induced penetration across the interface leads to highly localized strain concentrations and permanent material damage. These observations suggest that residual strain is primarily generated during phases of maximum deformation, including (i) the expansion stage near R_max_, where hoop stretch is maximized, and (ii) asymmetric collapse events that induce strong interfacial deformation and localized material failure.

Our experimental observations can provide insight into other soft matter interface scenarios that could feature interfacial cavitation, such as recent reports on subsurface gray matter cellular damage in human cortical brain tissue samples exposed to blasts [63, 64]. The study discussed in S. Shively et al. [65] highlights the formation of astroglial scarring at specific interfaces within the brain, particularly at boundaries between different types of tissue and fluid compartments with similar impedance differences as observed in our study. Thus perhaps, the damage patterns, i.e., residual stresses and strains, observed in our gelatin hydrogels may provide some physical insight into a potential cavitation-based damage mechanism caused by blast exposures that contribute to damage at interfaces within biological tissues at extremely high rates, which would be consistent with the length and time scales described [65] but yet to be proven objectively.

## 4 Conclusion

In this paper, we developed an experimental framework to quantify inertial cavitation dynamics near a compliant gel–water interface. Using ultrafast imaging and our recently developed STAQ-DIC image tracking method, we captured full-field deformation, strain localization, and jet formation across a wide range of stand-off distances.

By systematically varying the non-dimensional stand-off distance *γ* = *d/R*_max_, we identified four distinct cavitation regimes including (i) bulk-like symmetric oscillations (*γ >* 0.62), (ii) interface-contact-induced asymmetry (0 *< γ <* 0.62), (iii) interface penetration (−0.62 *< γ <* 0), and (iv) fully confined cavitation within the gel (*γ <* −0.62). These regimes demonstrate that geometric proximity to an interface is a primary parameter governing cavitation morphology, jet directionality, and deformation patterns. By performing full-field deformation measurements, we measured complex subsurface kinematics, including stagnation points, vortex-like structures, and shear-wave-dominated deformation fields. These features highlight the strongly coupled fluid–structure interaction that arises when cavitation occurs near compliant interfaces, which cannot be captured by traditional surface-based or spherically symmetric models. Compared to previous literature studies where only positive values of stand-off distances have been investigated [23, 24], we also studied different negative stand-off distance cases in this paper. Importantly, we show that the critical stand-off distance for interface penetration is largely insensitive to gel stiffness across the range tested. This suggests that the governing mechanism extends beyond elastic stiffness and is instead controlled by large-deformation processes and potentially dynamic fracture.

The experiments demonstrate that even modest compliance or material property mismatches at an interface can fundamentally alter the cavitation behavior. The results highlight the need for interface-aware modeling in cavitation-driven processes within engineered hydrogels, biological tissues, and soft material systems. Beyond advancing the understanding of fluid–structure interaction during inertial cavitation, the present framework provides a quantitative foundation for predicting interface stresses and designing future applications in therapeutic ultrasound, cavitation-based material processing, and soft tissue biomechanics.

## Limitations and Future Work

Although the experimental framework developed in this study provides new insight into cavitation near soft interfaces, several limitations remain. First, the present measurements are inherently two-dimensional and capture only a single speckle plane. Out-of-plane motion and three-dimensional bubble asymmetries, particularly during jetting or toroidal collapse, may not be fully resolved. Extending the approach to volumetric STAQ-DIC or tomographic imaging would enable direct quantification of the full 3D deformation field and improve model calibration.

Second, the stress fields reported in this work are inferred using constitutive models calibrated from independent experiments and do not explicitly account for material damage or failure. However, the large, localized deformations observed near the interface suggest that damage, microstructural failure, and dynamic fracture likely play an important role. Incorporating damage mechanics and fracture models into both experimental interpretation and theoretical frameworks will be essential to fully understand cavitation-induced material failure.

Third, all experiments were conducted under single cavitation events. In many practical applications, such as histotripsy, lithotripsy, and repetitive laser pulsing, materials are subjected to repeated cavitation cycles and multi-bubble interactions. These effects may lead to cumulative damage, altered material properties, and complex bubble–bubble coupling [66]. Future studies will investigate multi-cycle cavitation and collective bubble dynamics to better represent application-relevant conditions.

Finally, while the present work identifies key phenomenological regimes governed by the stand-off distance *γ*, a predictive theoretical or computational model that captures the coupled effects of interface proximity, material nonlinearity, and dynamic fracture is still lacking. Developing such models, validated against the full-field experimental data presented here, represents an important direction for future research.

## Acknowledgment

We gratefully acknowledge support from the US Office of Naval Research under PANTHER award numbers N000142112044 and N000142212828 through Dr. Timothy Bentley. J.Y. and M.R.J. acknowledge the support provided by the U.S. National Science Foundation (NSF) under Grant No. CMMI-2232427, CMMI-2232428, respectively. M.R.J. acknowledges support from the US Department of Defense under the DEPSCoR program Award No. FA9550-23-1-0485 through Dr. Timothy Bentley. J.Y. acknowledges the start-up fund from the University of Texas at Austin and the Haythornthwaite Foundation for the Research Initiation Grant awarded by the Applied Mechanics Division of the American Society of Mechanical Engineers (ASME).

## Declarations

### Ethical Approval

This study complies with chemical and biological safety requirements under approved protocol at the University of Wisconsin-Madison and the IBC-2025-00178 at the University of Texas at Austin.

### Consent to Participate and Consent to Publish

All authors consent to participate in this study and to the publication of this manuscript.

### Author Contributions

Jin Yang and Alexander McGhee conceived the research, designed and conducted the experiments. Jin Yang and Alexander McGhee mentored Griffin Radtke to analyze the bubble penetration behavior. Jin Yang and Zixiang Tong performed DIC post-processing. Jin Yang, Christian Franck, and Alexander McGhee prepared the original draft of the manuscript. All authors reviewed and edited the manuscript and approved the final version.

### Availability of Data and Materials

The data that support the findings of this study are available from the corresponding author upon reasonable request.

## Footnotes

1 The von Mises strain is defined using a similar mathematical expression as von Mises stress but using infinitesimal strain components

## References

1. Michel Versluis, Barbara Schmitz, Anna von der Heydt, and Detlef Lohse. How snapping shrimp snap: through cavitating bubbles. Science, 289(5487):2114–2117, 2000.

2. Detlef Lohse, Barbara Schmitz, and Michel Versluis. Snapping shrimp make flashing bubbles. Nature, 413(6855):477–478, 2001.

3. Jules W Lindau, David A Boger, Richard B Medvitz, and Robert F Kunz. Propeller cavitation breakdown analysis. Journal of Fluids Engineering, 127:995–1002, 2005.

4. RH Cole. Underwater explosions, 1948.

5. Christopher E Brennen. Cavitation and bubble dynamics. Cambridge university press, 2014.

6. Benjamin Dollet, Philippe Marmottant, and Valeria Garbin. Bubble dynamics in soft and biological matter. Annual Review of Fluid Mechanics, 51(1):331–355, 2019.

7. Christopher W Barney, Carey E Dougan, Kelly R McLeod, Amir Kazemi-Moridani, Yue Zheng, Ziyu Ye, Sacchita Tiwari, Ipek Sacligil, Robert A Riggleman, Shengqiang Cai, et al. Cavitation in soft matter. Proceedings of the National Academy of Sciences, 117(17):9157–9165, 2020.

8. Chunghwan Kim, Won June Choi, Yisha Ng, and Wonmo Kang. Mechanically induced cavitation in biological systems. Life, 11(6):546, 2021.

9. Lord Rayleigh. On the Pressure developed in a Liquid during the Collapse of a Spherical Cavity. Philosophical Magazine, 6:94–98, 1917.

10. M. S. Plesset. The Dynamics of Cavitation Bubbles. Journal of Applied Mechanics, 16:277–282, 1949.

11. C. Herring. Theory of the Pulsations of the Gas Bubble Produced by an Underwater Explosion. Technical report, OSRD Report No. 236, 1941.

12. F R Gilmore. The growth or collapse of a spherical bubble in a viscous compressible liquid. Technical report, Office of Naval Research, Pasadena, CA, 1952.

13. Leon Trilling. The Collapse and Rebound of a Gas Bubble. Journal of Applied Physics, 23:14–17, 1952.

14. Joseph B. Keller and Ignace I. Kolodner. Damping of Underwater Explosion Bubble Oscillations. Journal of Applied Physics, 27:1152–1161, 1956.

15. Robert Hickling and Milton S. Plesset. Collapse and Rebound of a Spherical Bubble in Water. Physics of Fluids, 7:1–7, 1964.

16. H. G. Flynn. Cavitation dynamics. i. a mathematical formulation. Journal of the Acoustical Society of America, 57:1379–1396, 1975.

17. Joseph B Keller and Michael Miksis. Bubble oscillations of large amplitude. The Journal of the Acoustical Society of America, 68(2):628–633, 1980.

18. Shigeo Fujikawa and Teruaki Akamatsu. Effects of the non-equilibrium condensation of vapour on the pressure wave produced by the collapse of a bubble in a liquid. Journal of Fluid Mechanics, 97:481–512, 1980.

19. R Gaudron, MT Warnez, and E Johnsen. Bubble dynamics in a viscoelastic medium with nonlinear elasticity. Journal of Fluid Mechanics, 766:54–75, 2015.

20. Jonathan B Estrada, Carlos Barajas, David L Henann, Eric Johnsen, and Christian Franck. High strain-rate soft material characterization via inertial cavitation. Journal of the Mechanics and Physics of Solids, 112:291–317, March 2018.

21. Anastasia Tzoumaka, Jin Yang, Selda Buyukozturk, Christian Franck, and David L Henann. Modeling high strain-rate microcavitation in soft materials: the role of material behavior in bubble dynamics. Soft Matter, 19(21):3895–3909, 2023.

22. John R Blake and DC Gibson. Cavitation bubbles near boundaries. Annual Review of Fluid Mechanics, 19(1):99–123, 1987.

23. Emil-Alexandru Brujan, Kester Nahen, Peter Schmidt, and Alfred Vogel. Dynamics of laser-induced cavitation bubbles near an elastic boundary. Journal of Fluid Mechanics, 433:251–281, 2001.

24. Emil-Alexandru Brujan, Kester Nahen, Peter Schmidt, and Alfred Vogel. Dynamics of laser-induced cavitation bubbles near elastic boundaries: influence of the elastic modulus. Journal of Fluid Mechanics, 433:283–314, 2001.

25. Shucheng Pan, Stefan Adami, Xiangyu Hu, and Nikolaus A Adams. Phenomenology of bubble-collapse-driven penetration of biomaterial-surrogate liquid-liquid interfaces. Physical Review Fluids, 3(11):114005, 2018.

26. Uroš Orthaber, Jure Zevnik, Matevž Dular, et al. Cavitation bubble collapse in a vicinity of a liquid-liquid interface–basic research into emulsification process. Ultrasonics Sonochemistry, 68:105224, 2020.

27. Thomas Henzel, Japinder Nijjer, S Chockalingam, Hares Wahdat, Alfred J Crosby, Jing Yan, and Tal Cohen. Interfacial cavitation. PNAS Nexus, 1(4):pgac217, 2022.

28. Pooya Movahed, Wayne Kreider, Adam D. Maxwell, Shelby B. Hutchens, and Jonathan B. Freund. Cavitation-induced damage of soft materials by focused ultrasound bursts: A fracture-based bubble dynamics model. J. Acoust. Soc. Am., 140(2):1374–1386, 2016.

29. Shabnam Raayai-Ardakani, Darla Rachelle Earl, and Tal Cohen. The intimate relationship between cavitation and fracture. Soft Matter, 15(25):4999–5005, 2019.

30. Matt P Milner and Shelby B Hutchens. Dynamic fracture of expanding cavities in nonlinear soft solids. Journal of Applied Mechanics, 88(8):081008, 2021.

31. Matt P Milner and Shelby B Hutchens. Multi-crack formation in soft solids during high rate cavity expansion. Mechanics of Materials, 154:103741, 2021.

32. Jingtian Kang, Changguo Wang, and Shengqiang Cai. Cavitation to fracture transition in a soft solid. Soft Matter, 13(37):6372–6376, 2017.

33. Martin Oliver Steinhauser and Mischa Schmidt. Destruction of cancer cells by laser-induced shock waves: recent developments in experimental treatments and multiscale computer simulations. Soft Matter, 10(27):4778–4788, 2014.

34. Jonathan A Kopechek, Eun-Joo Park, Yong-Zhi Zhang, Natalia I Vykhodtseva, Nathan J McDan-nold, and Tyrone M Porter. Cavitation-enhanced mr-guided focused ultrasound ablation of rabbit tumors in vivo using phase shift nanoemulsions. Physics in Medicine & Biology, 59(13):3465, 2014.

35. Constantin C Coussios and Ronald A Roy. Applications of acoustics and cavitation to noninvasive therapy and drug delivery. Annual Review of Fluid Mechanics, 40:395–420, 2008.

36. Eleanor Stride and Constantin Coussios. Nucleation, mapping and control of cavitation for drug delivery. Nature Reviews Physics, 1(8):495–509, 2019.

37. Zoltan Z Nagy. New technology update: femtosecond laser in cataract surgery. Clinical Ophthalmology, pages 1157–1167, 2014.

38. Peter Johansen. Mechanical heart valve cavitation. Expert Review of Medical Devices, 1(1):95–104, 2004.

39. Wayne Kreider, Lawrence A Crum, Michael R Bailey, and Oleg A Sapozhnikov. Observations of the collapses and rebounds of millimeter-sized lithotripsy bubbles. The Journal of the Acoustical Society of America, 130(5):3531–3540, 2011.

40. Alfred Vogel, Werner Hentschel, Joachim Holzfuss, and Werner Lauterborn. Cavitation bubble dynamics and acoustic transient generation in ocular surgery with pulsed neodymium: YAG lasers. Ophthalmology, 93(10):1259–1269, 1986.

41. Ji Lang, Rungun Nathan, Dong Zhou, Xuewei Zhang, Bo Li, and Qianhong Wu. Cavitation causes brain injury. Physics of Fluids, 33(3), 2021.

42. Fuad Hasan, KAH Al Mahmud, Md Ishak Khan, Sandeep Patil, Brian H Dennis, and Ashfaq Adnan. Cavitation induced damage in soft biomaterials. Multiscale Science and Engineering, 3(1):67–87, 2021.

43. Jonathan B Estrada, Harry C Cramer III, Mark T Scimone, Selda Buyukozturk, and Christian Franck. Neural cell injury pathology due to high-rate mechanical loading. Brain Multiphysics, 2:100034, 2021.

44. J Yang, H C Cramer, III, and C Franck. Extracting non-linear viscoelastic material properties from violently-collapsing cavitation bubbles. Extreme Mechanics Letters, 2020.

45. Jin Yang, Harry C Cramer, Elizabeth C Bremer, Selda Buyukozturk, Yue Yin, and Christian Franck. Mechanical characterization of agarose hydrogels and their inherent dynamic instabilities at ballistic to ultra-high strain-rates via inertial microcavitation. Extreme Mechanics Letters, 51:101572, February 2022.

46. EC Bremer-Sai, J Yang, A McGhee, and C Franck. Ballistic and blast-relevant, high-rate material properties of physically and chemically crosslinked hydrogels. Experimental Mechanics, 64(4):587–592, 2024.

47. Michael A Sutton, Jean Jose Orteu, and Hubert Schreier. Image Correlation for Shape, Motion and Deformation Measurements: Basic Concepts, Theory and Applications. Springer Science & Business Media, April 2009.

48. J Yang, V Rubino, Z Ma, J Tao, Y Yin, A McGhee, W Pan, and C Franck. Spatiotemporally adaptive quadtree mesh (staq) digital image correlation for resolving large deformations around complex geometries and discontinuities. Experimental Mechanics, 62(7):1191–1215, 2022.

49. A McGhee, A Bennett, P Ifju, GW Sawyer, and TE Angelini. Full-field deformation measurements in liquid-like-solid granular microgel using digital image correlation. Experimental Mechanics, 58:137–149, 2018.

50. AJ McGhee, EO McGhee, JE Famiglietti, and KD Schulze. Dynamic subsurface deformation and strain of soft hydrogel interfaces using an embedded speckle pattern with 2D digital image correlation. Experimental Mechanics, 61:1017–1027, 2021.

51. A McGhee, J Yang, EC Bremer, Z Xu, HC Cramer III, JB Estrada, DL Henann, and CJEM Franck. High-speed, full-field deformation measurements near inertial microcavitation bubbles inside viscoelastic hydrogels. Experimental Mechanics, 63(1):63–78, 2023.

52. Lehu Bu, Zhao-Bang Hou, Sophie Polidoro, and Jin Yang. High-speed, full-field measurement of large deformations near needle-induced cavitation bubbles within biological soft materials. In SEM Annual Conference and Exposition on Experimental and Applied Mechanics, pages 115–125. Springer, 2024.

53. Fabian Reuter, Qingyun Zeng, and Claus-Dieter Ohl. The rayleigh prolongation factor at small bubble to wall stand-off distances. Journal of Fluid Mechanics, 944:A11, 2022.

54. AB Sieber, DB Preso, and M Farhat. Cavitation bubble dynamics and microjet atomization near tissue-mimicking materials. Physics of Fluids, 35(2), 2023.

55. Gabriel Taubin. Estimation of planar curves, surfaces, and nonplanar space curves defined by implicit equations with applications to edge and range image segmentation. IEEE Transactions on Pattern Analysis & Machine Intelligence, 13(11):1115–1138, 1991.

56. J Yang, A Tzoumaka, K Murakami, E Johnsen, and others. Predicting complex non-spherical instability shapes of inertial cavitation bubbles in viscoelastic soft matter. arXiv preprint arXiv, 2021.

57. Xiaoxu Zhong, Javad Eshraghi, Pavlos Vlachos, Sadegh Dabiri, and Arezoo M Ardekani. A model for a laser-induced cavitation bubble. International Journal of Multiphase Flow, 132:103433, 2020.

58. Xiao-Xuan Liang, Norbert Linz, Sebastian Freidank, Günther Paltauf, and Alfred Vogel. Comprehensive analysis of spherical bubble oscillations and shock wave emission in laser-induced cavitation. Journal of Fluid Mechanics, 940:A5, 2022.

59. AM Zhang, Shi-Min Li, Pu Cui, Shuai Li, and Yun-Long Liu. A unified theory for bubble dynamics. Physics of Fluids, 35(3), 2023.

60. Jin Yang, Yue Yin, Harry C Cramer, and Christian Franck. The penetration dynamics of a violent cavitation bubble through a hydrogel–water interface. In Challenges in Mechanics of Time Dependent Materials, Mechanics of Biological Systems and Materials & Micro-and Nanomechanics, Volume 2: Proceedings of the 2021 Annual Conference & Exposition on Experimental and Applied Mechanics, pages 65–71. Springer, 2022.

61. Justin C. Luo, Herman Ching, Bryce G. Wilson, Ali Mohraz, Elliot L. Botvinick, and Vasan Venugopalan. Laser cavitation rheology for measurement of elastic moduli and failure strain within hydrogels. Scientific Reports, 10(1), AUG 4 2020.

62. Zhiren Zhu, Sawyer Remillard, Bachir A Abeid, Danila Frolkin, Spencer H Bryngelson, Jin Yang, Mauro Rodriguez, and Jonathan B Estrada. Parsimonious inertial cavitation rheometry via bubble collapse time. Soft Matter, 21(34):6717–6734, 2025.

63. Christian Franck. Microcavitation: the key to modeling blast traumatic brain injury? Concussion, 2(3):CNC47, 2017.

64. Alice Lux Fawzi and Christian Franck. Beyond symptomatic diagnosis of mild traumatic brain injury. Concussion, 8(2), 2023.

65. Sharon Baughman Shively, Iren Horkayne-Szakaly, Robert V Jones, James P Kelly, Regina C Armstrong, and Daniel P Perl. Characterisation of interface astroglial scarring in the human brain after blast exposure: a post-mortem case series. The Lancet Neurology, 15(9):944–953, 2016.

66. Jin Yang, Alexander McGhee, Zixiang Tong, Lehu Bu, Sicong Wang, Griffin Radtke, Mauro Rodriguez, and Christian Franck. Spatiotemporally-resolved kinematic and stress measurements of interfacial cavitation in soft matter via dic. In Computer Vision & Laser Vibrometry*, Vol.* 6, pages 1–12. River Publishers, 2026.

