## Supplemental PDF 1 for "Inertial interface cavitation creates complex, flow-like structures within a soft solid"

**Noname manuscript No.**
(will be inserted by the editor)

### Supplementary Material of "Inertial interface cavitation creates complex, flow-like structures within a soft solid"

Jin Yang [1,2,†*] · Alexander McGhee [3†] · Zixiang Tong [1] ·
Griffin Radtke [4] · Mauro Rodriguez Jr. [5] · Christian Franck [6‡]

#### S1 STAQ-DIC tracked full-field kinematic and stress fields for nondimensional stand-off distance $\gamma = 2.19$

Analysis of strain and stress fields during laser-induced inertial cavitation near the gel-water interface with a nondimensional stand-off distance $\gamma = 2.19$ are summarized in Fig. S1.

For each time frame, narrow vertical and horizontal slices extracted from the full-field DIC results were concatenated in time to construct the kymographs. The inset schematic on the left illustrates the experimental geometry, where $d$ denotes the distance from the bubble center to the gel–water interface and $h$ represents the depth coordinate within the gelatin hydrogel. The results show that, for $\gamma = 2.19$, the cavitation bubble primarily induces localized elastic deformation near the interface during expansion and collapse, followed by the propagation of shear-dominated stress waves into the bulk gel after the first collapse event.

[1]Department of Aerospace Engineering and Engineering Mechanics, The University of Texas at Austin, Austin, TX, USA 78712

[2]Texas Materials Institute, The University of Texas at Austin, Austin, TX, USA 78712

[3]Department of Biomedical Engineering, The University of Arizona, Tucson, AZ, USA 85721

[4]John A. Paulson School of Engineering and Applied Sciences, Harvard University, Cambridge, MA, USA 02138

[5]School of Engineering, Brown University, Providence, RI, USA 02912

[6]Department of Mechanical Engineering, University of Wisconsin-Madison, Madison, WI USA 53706

[†]J.Y. and A.M. contributed equally to this work.

[*]

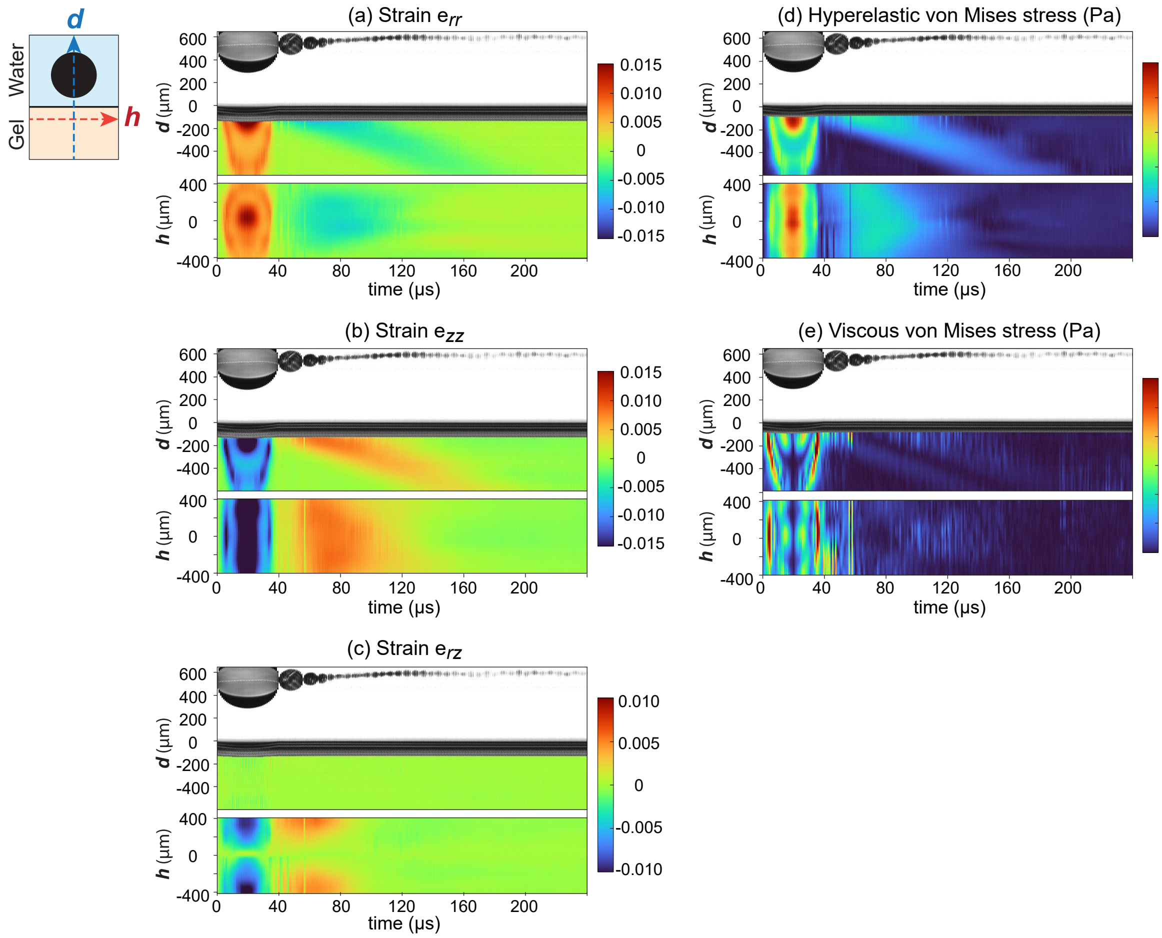


Fig. S1: Spatiotemporal evolution of experimentally measured strain and stress fields during laser-induced inertial cavitation near the gel–water interface for a non-dimensional stand-off distance of $\gamma = 2.19$. (a–c) Kymographs of the DIC-measured strain components $e_{rr}$, $e_{zz}$, and $e_{rz}$, respectively. (d-e) Hyperelastic von Mises stress estimated using the neo-Hookean Kelvin–Voigt constitutive framework for hyperelastic and viscous components, respectively.

#### S2 STAQ-DIC tracked full-field kinematic and stress fields for nondimensional stand-off distance $\gamma = 0.38$

Analysis of strain and stress fields during laser-induced inertial cavitation near the gel-water interface with a nondimensional stand-off distance $\gamma = 0.38$ are summarized in Fig. S2.

To improve tracking robustness throughout the cavitation cycle, different DIC tracking strategies were adopted during post-processing. During the early stages of bubble evolution (from $t = 0$ to the vertical dashed line in the kymographs), incremental frame-to-frame tracking was used to accurately capture the rapidly evolving deformation field. After this time point, cumulative tracking relative to the undeformed reference configuration was employed to quantify the long-term residual deformation and stress evolution within the hydrogel. The viscous von Mises stress fields shown in Fig. S2(e) were computed entirely using incremental tracking results because viscous stresses depend directly on instantaneous deformation-rate and velocity-gradient measurements.

For each DIC frame, narrow vertical and horizontal slices extracted from the full-field DIC measurements were concatenated in time to construct the kymographs. The inset schematic on the left illustrates the experimental geometry, where $d$ denotes the distance from the bubble center to the gel–water interface, while $h_1$ and $h_2$ correspond to two different depths within the gelatin hydrogel.

Compared with the $\gamma = 2.19$ case in Fig. S1, the smaller stand-off distance produces substantially larger deformation and stress concentrations near the interface due to stronger bubble–interface interactions.

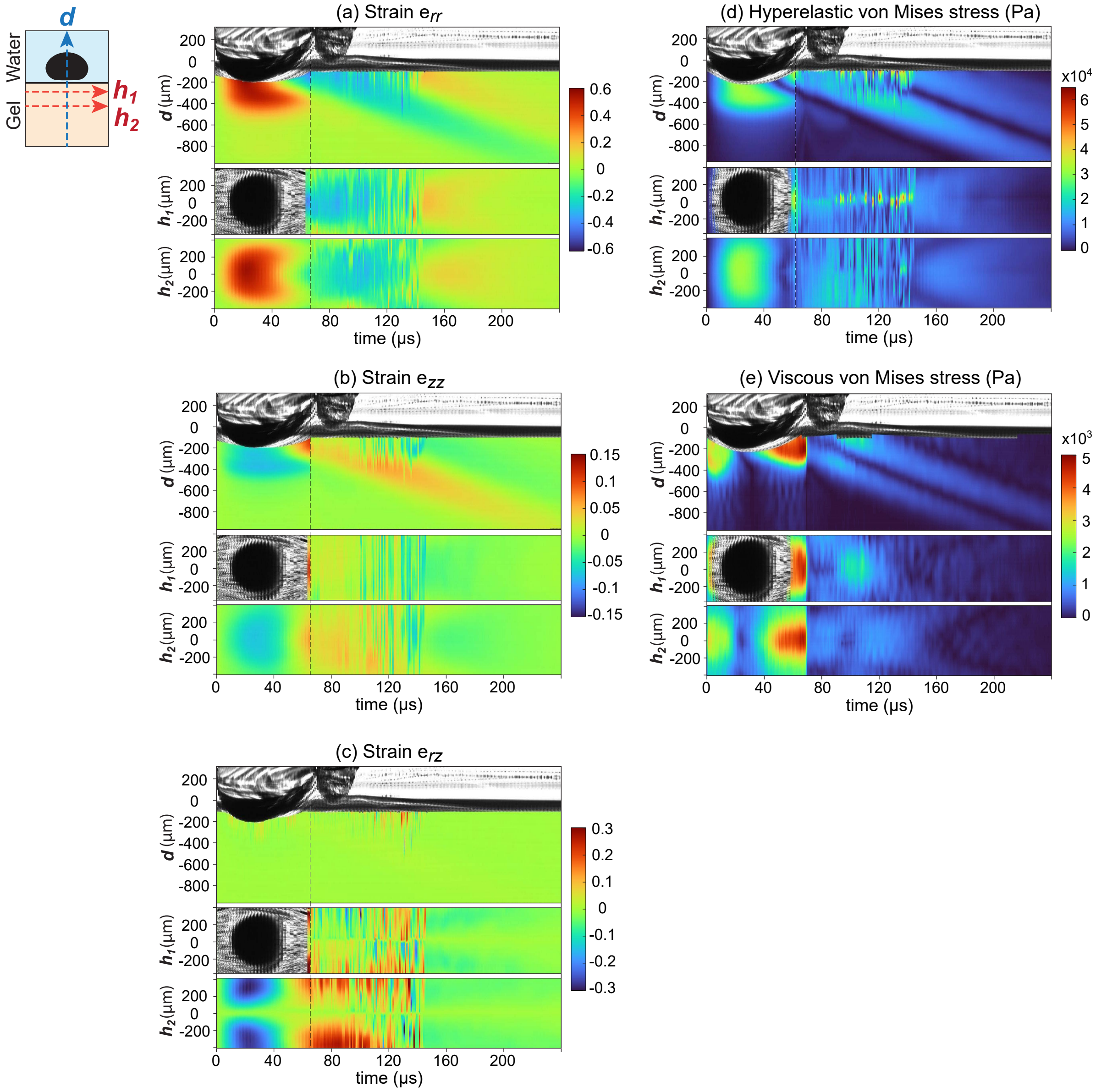


Fig. S2: Spatiotemporal evolution of experimentally measured strain and stress fields during laser-induced inertial cavitation near the gel–water interface for a non-dimensional stand-off distance of $\gamma = 0.38$. (a–c) Kymographs of the DIC-measured strain components $e_{rr}$, $e_{zz}$, and $e_{rz}$, respectively. (d-e) Hyperelastic von Mises stress estimated using the neo-Hookean Kelvin–Voigt constitutive framework for hyperelastic and viscous components, respectively.

#### S3 STAQ-DIC tracked full-field kinematic and stress fields for nondimensional stand-off distance $\gamma = -0.56$

Analysis of strain and stress fields during laser-induced inertial cavitation near the gel-water interface with a nondimensional stand-off distance $\gamma = -0.56$ are summarized in Fig. S3.

To improve tracking robustness throughout the cavitation cycle, different DIC tracking strategies were adopted during post-processing. During the early stages of bubble evolution (from $t = 0$ to the vertical dashed line in the kymographs), incremental frame-to-frame tracking was used to accurately capture the rapidly evolving deformation field. After this time point, cumulative tracking relative to the undeformed reference configuration was employed to quantify the long-term residual deformation and stress evolution within the hydrogel. The viscous von Mises stress fields shown in Fig. S3(e) were computed entirely using incremental tracking results because viscous stresses depend directly on instantaneous deformation-rate and velocity-gradient measurements.

For each DIC frame, narrow vertical and horizontal slices extracted from the full-field DIC measurements were concatenated in time to construct the kymographs. The inset schematic on the left illustrates the experimental geometry, where $d$ denotes the distance from the bubble center to the gel–water interface and $h$ represents the depth coordinate within the gelatin hydrogel.

For $\gamma = -0.56$, the cavitation bubble nucleates within the gel and subsequently penetrates through the gel–water interface into the water phase. Compared with the cases shown in Figs. S1-S2, this regime generates significantly larger and more localized deformation fields near the interface. The strain kymographs reveal strong asymmetric deformation during bubble expansion and collapse, accompanied by substantial shear strain localization following interface penetration. Persistent residual strain and stress signatures indicate substantial irreversible deformation and localized material damage associated with this cavitation regime.

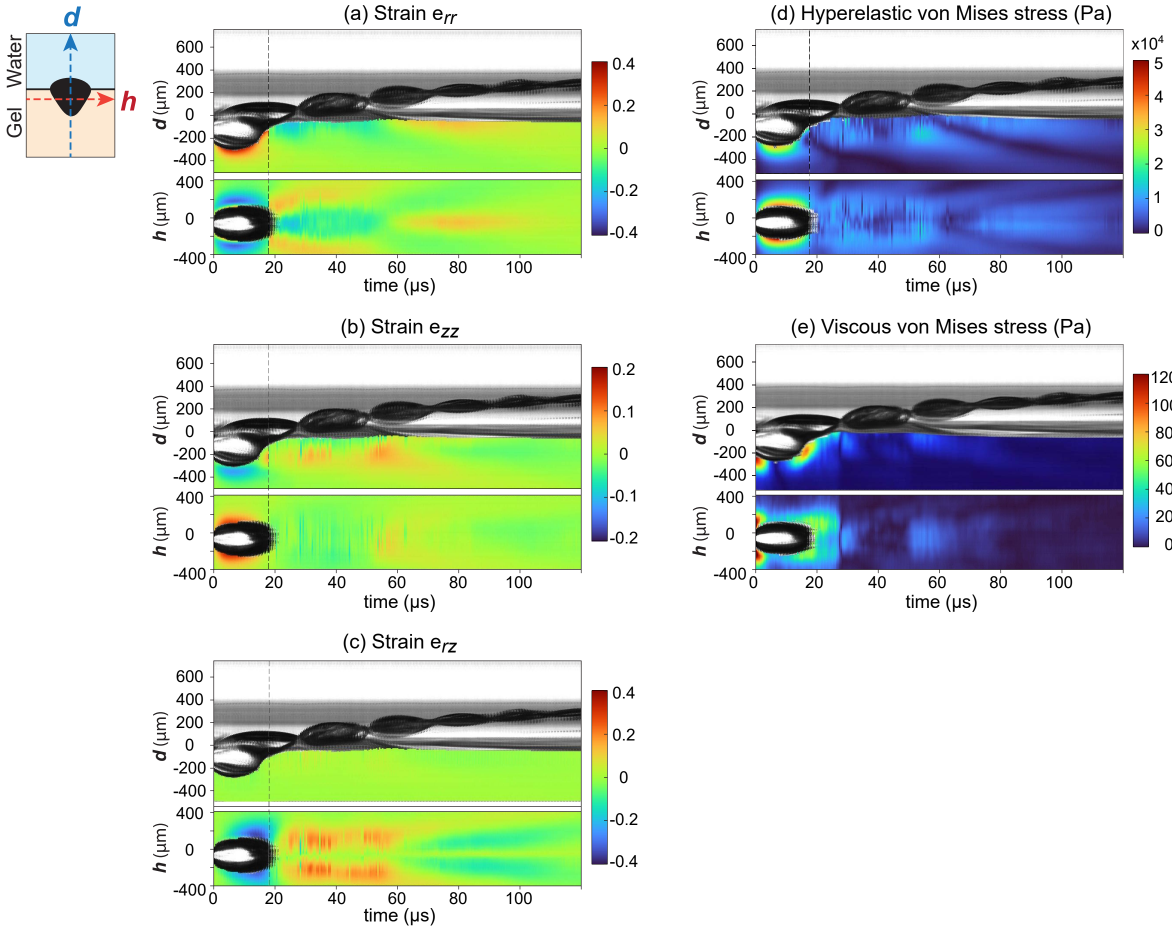


Fig. S3: Spatiotemporal evolution of experimentally measured strain and stress fields during laser-induced inertial cavitation near the gel–water interface for a non-dimensional stand-off distance of $\gamma = -0.56$. (a–c) Kymographs of the DIC-measured strain components $e_{rr}$, $e_{zz}$, and $e_{rz}$, respectively. (d-e) Hyperelastic von Mises stress estimated using the neo-Hookean Kelvin–Voigt constitutive framework for hyperelastic and viscous components, respectively.

#### S4 STAQ-DIC tracked full-field kinematic and stress fields for nondimensional stand-off distance $\gamma = -3.16$

Analysis of strain and stress fields during laser-induced inertial cavitation near the gel-water interface with a nondimensional stand-off distance $\gamma = -3.16$ are summarized in Fig. S4.

To improve tracking robustness throughout the cavitation cycle, different DIC tracking strategies were adopted during post-processing. During the early stages of bubble evolution (from $t = 0$ to the vertical dashed line in the kymographs), incremental frame-to-frame tracking was used to accurately capture the rapidly evolving deformation field. After this time point, cumulative tracking relative to the undeformed reference configuration was employed to quantify the long-term residual deformation and stress evolution within the hydrogel. The viscous von Mises stress fields shown in Fig. S4(e) were computed entirely using incremental tracking results because viscous stresses depend directly on instantaneous deformation-rate and velocity-gradient measurements.

For each DIC frame, narrow vertical and horizontal slices extracted from the full-field DIC measurements were concatenated in time to construct the kymographs. The inset schematic on the left illustrates the experimental geometry, where $d$ denotes the distance from the bubble center to the gel–water interface, while $h_1$ and $h_2$ correspond to two different depths within the gelatin hydrogel.

For $\gamma = -3.16$, the cavitation bubble nucleates and remains entirely within the gelatin hydrogel, resulting in deformation dynamics that closely resemble bulk cavitation behavior. Compared with the more interfacially dominated cases shown in Figs. S2-S3, the strain and stress fields exhibit substantially greater spatial symmetry about the bubble center. The strain kymographs reveal predominantly radial tensile and compressive deformation during the expansion and collapse phases, while the shear strain component $\varepsilon_{rz}$ remains relatively localized and weak except during the later collapse and rebound stages.

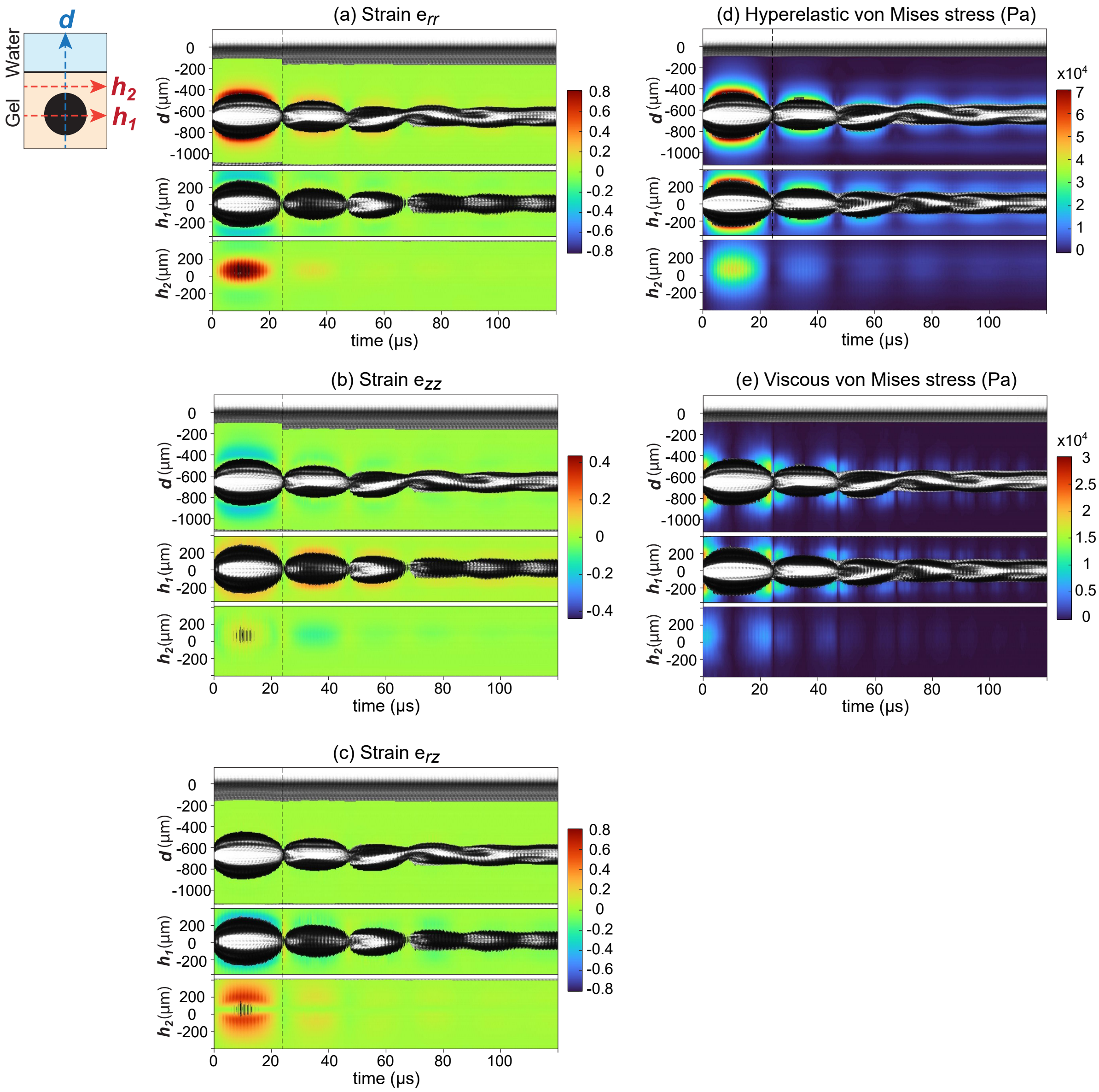


Fig. S4: Spatiotemporal evolution of experimentally measured strain and stress fields during laser-induced inertial cavitation near the gel–water interface for a non-dimensional stand-off distance of $\gamma = -3.16$. (a–c) Kymographs of the DIC-measured strain components $e_{rr}$, $e_{zz}$, and $e_{rz}$, respectively. (d-e) Hyperelastic von Mises stress estimated using the neo-Hookean Kelvin–Voigt constitutive framework for hyperelastic and viscous components, respectively.
